# Crucial role for iron metabolism in mediating influenza A virus infection and associated disease

**DOI:** 10.1101/2025.03.20.644262

**Authors:** Amber L. Pillar, Katie Daly, Henry M. Gomez, Ama-Tawiah Essilfie, Kai Sen Tan, Jing Liu, Anand Kumar Andiappan, Alexandra C. Brown, Richard Y. Kim, Kristy Nichol, Chantal Donovan, Greg J. Anderson, Andreas Suhrbier, David Frazer, Elizabeth A Milward, Vincent T Chow, Mookkan Prabakaran, De Yun Wang, Philip M. Hansbro, David W. Reid, Alan C. Y. Hsu, Peter A. B. Wark, Jay C. Horvat, Jemma R. Mayall

## Abstract

**Rationale and Objectives:** Iron availability and metabolism are important in the pathogenesis of bacterial infections. More recently, links have been reported between iron and the severity of viral infections. In this study, we characterize a crucial relationship between iron metabolism and IAV infection and disease.

**Methods:** Iron-related gene expression was assessed in human airway epithelial cells (AEC) infected with IAV. AECs were cultured with ferric iron, iron-loaded transferrin, or iron chelator, deferoxamine (DFO), prior to infection with IAV. Mice were placed on a high iron diet for 8 weeks prior to infection with IAV or treated with anti-transferrin receptor-1 (TFR1) antibody during IAV infection. The effects of iron modulation and depletion of TFR1-mediated responses on IAV infection were assessed.

**Measurements and main results:** Iron-related gene expression and metabolism are altered systemically and in lung tissues and AECs during IAV infections. Increasing iron availability increases viral titer in AECs, while DFO protects against iron-induced increased susceptibility to infection. Increasing systemic iron loading, which increases iron levels in the lung, increases viral titer, proinflammatory responses, airway inflammation, and worsens IAV-induced disease in terms of lung function and weight loss *in vivo*. Inhibition of TFR1 protects against IAV-induced disease *in vivo*.

**Conclusion:** IAV infections remain a major threat to human health and global economies. Strategies that boost protective, or reduce pathogenic, host responses may provide broadly effective, long-term therapeutic options. We have identified a key role for iron metabolism in modifying host responses to IAV that can be harnessed to protect against disease.

**Key Messages:**

- Iron metabolism is altered systemically, and in airway epithelial cells and lung tissues, during IAV infection.
- Increased iron availability increases viral titer both *in vitro* and *in vivo*
- Systemic iron loading worsens IAV-induced inflammation and disease outcomes *in vivo,* highlighting iron as a crucial factor for modulating IAV infections and disease.
- Host epithelial cells and lung tissues reduce *TFRC* gene expression, whilst the number and proportion of TFR1^hi^ expressing cells increase, in response to IAV infection. Neutralising TFR1 protects against IAV-induced disease *in vivo,* highlighting TFR1 as a potential therapeutic target for IAV infections.

## INTRODUCTION

Iron is essential for cellular respiration, the function of many enzymes, and cell proliferation in virtually all living organisms. Iron is transported in the blood to organs primarily *via* transferrin (Tf) and Tf-Fe^3+^ (holo-Tf) is endocytosed into all nucleated cells by transferrin receptor-1 (TFR1) (1). Cytosolic iron is then either stored in ferritin, which is upregulated in response to increasing intracellular iron levels, incorporated into enzymes for use in various metabolic processes, or released back into the extracellular space *via* the only known iron exporter, ferroportin (encoded by *SLC40A1*) (2–4). Macrophages play a central role in the regulation of iron homeostasis in tissues through the storage of Tf-derived iron and recycling of heme-iron from senescent RBCs (2).

The interactions between airway epithelial cells and immune cells play a critical role in mucosal iron homeostasis, protecting lung cells from iron deficiency-induced loss of function and iron-induced oxidative stress (2). It is well established that iron availability and metabolism play key roles in the pathogenesis of both extracellular and intracellular bacterial infections, largely due to iron being required for bacterial metabolism (2). Importantly, host responses have evolved to limit iron availability to bacteria, particularly at mucosal sites, by altering iron uptake, storage, and export. The handling and storage of iron by macrophages and epithelial cells play particularly important roles in and in sequestering iron from invading bacterial pathogens (2).

Evidence shows that iron availability also plays a critical role in viral infections. Increased iron availability has been shown to increase viral replication and susceptibility to infections for HIV-1, hepatitis B and C viruses, human cytomegalovirus, vaccinia virus, and herpes simplex virus-1 (5–9). Recently, strong correlations have been identified between iron levels and SARS-CoV-2-associated morbidity and mortality (10–13) and *in vitro* studies have identified ferroptosis as a response to swine influenza infection that enhances viral replication (14). However, studies investigating the association between iron status and respiratory viral infection are predominantly based on clinical associations and/or *in vitro* studies (10–12). There are few comprehensive functional studies investigating the role of iron levels and modulation at the cellular and organism level. Importantly, whether iron availability affects susceptibility to respiratory viral infections or associated disease, and whether systemic, airways or cellular iron levels can be manipulated therapeutically to treat infection-induced disease, has not been fully characterised for respiratory viruses.

In this study, we investigated IAV-induced changes to iron metabolism, and the impact of modulating iron levels and metabolism in human epithelial cell and murine models, on IAV infection and IAV-induced disease, to better understand how iron metabolism affects, and may be therapeutically modulated to treat and prevent, IAV infection and associated disease.

## METHODS

Full details are provided in the supplementary materials.

### Study approvals

All experiments were conducted with approval of the Animal Care and Ethics Committee of the University of Newcastle, Australia (A-2021-135), Hunter New England Human Research Ethics Committee, Australia (05/08/10/3.09), Queensland Institute of Medical Research (QIMR) Berghofer Research Institute (P3778), National Healthcare Group Domain-Specific Board of Singapore (DSRB Ref: D/11/228), or Institutional Review Board of the National University of Singapore (IRB Ref: 13-509).

### Human AEC IAV infections

BCi-Ns1.1 cells (15) were stimulated with ferric (Fe^3+^) ammonium citrate (FAC), holo-Tf, or deferoxamine (DFO) at 10, 50 or 100μM for 24 hours prior to H1N1 infection (A/Auckland/1/2009; multiplicity of infection [MOI] of 0.5) and cells and cell supernatant collected at 2, 6, 12 and 24 hours post-infection (hpi; **Figure S1**) (16, 17). Primary nasal AECs grown at the air liquid interface (ALI) were infected with H1N1 (A/Singapore/G2-25.1/2014), H3N2 (A/Singapore/CDC-204/2014H3N2), influenza B virus (B/Singapore/G2-14.1/2014 (Victoria lineage)) or H5N1 (A/Indonesia/CDC-1031/2007) at MOI 0.1 and cells collected for RNA expression analysis at 24 hpi (18).

### Murine IAV infections

BALB/c mice were infected with H1N1 (A/Puerto Rico/8/1934; 7.5PFU) or sham-infected (19, 20). Groups of mice were fed a high iron diet (HID; 2% carbonyl iron) or control diet (CD) for 8 weeks prior to infection (2, 21), or administered anti-mouse TFR1 (αTFR1) or isotype control (Iso) antibody (2.5mg/kg) on days 0, 2 and 4 following infection. Lung function was measured and tissues collected for analyses at 3 and 5 days post-infection (dpi; **Figure S2**).

### Gene expression analyses

Gene expression in BCi-Ns1.1 cells (22) and murine lung and liver tissue (21, 23–27) was quantified via qPCR. Expression of iron metabolism related genes in primary AECs was separately quantified using RNA sequencing from human nasal AECs ALI cultures, infected with different strains and subtypes of influenza (MOI 0.1) (18). RNA isolation and sequencing were performed using extraction kits and library as previously described (18).

### Measurement of viral titer

Viral titre in cell supernatant and bronchoalveolar lavage fluid (BALF) was measured via plaque assay (19, 20), using a protocol adapted from (28).

### Measurement of cellular metabolism

Oxygen consumption rate and ATP production was measured in primary human airway epithelial cells (HBECs, LONZA) treated overnight with 100μM FAC using the Seahorse XF^e^96 Analyser and Cell Mito Stress assay kit.

### Measurement of non-heme iron levels, TFR1^hi^ cell numbers, lung function and airways inflammation

Iron accumulation in cells was assessed by enumerating the number of iron-loaded cells/mm^2^ in DAB-enhanced Perls-stained histological sections of mouse lung tissues (21, 23). Iron-loading cells were graded as low, intermediate and high. Non-heme iron levels were measured in murine lung and liver tissue by colorimetric assay (21, 23). The number and proportion of TFR1^hi^ immune and epithelial and endothelial cells were assessed in whole lung tissue using flow cytometry (21, 23). Lung function was measured at baseline and during airway hyperresponsiveness (AHR) using the FlexiVent system (21, 23–27, 29). Airway inflammation was also assessed by enumeration of leukocytes in BALF (21, 23–27, 29, 30).

### Statistics

Grubb’s outlier tests were performed and outliers removed. Two groups were compared using two-tailed t-tests, and multiple groups were compared using one-way analysis of variance (ANOVA) with either Fisher’s LSD post hoc tests. AHR and weight change data were assessed using two-way ANOVA with Fisher’s LSD post hoc tests. Differentially expressed genes from RNA sequencing were obtained following false discovery rate analysis, with gene expression presented as Log_2_FC. Analyses were performed using GraphPad Prism Software v10.0.2 (San Diego, California).

## RESULTS

### IAV infection alters iron metabolism in human AECs and murine lung tissue

To determine IAV infection affects iron metabolism in the lungs, we infected both human bronchial epithelial cells (BCi-Ns1.1 cell line) and BALB/c mice with IAV (H1N1) and assessed expression of genes for the factors that mediate cellular iron uptake, transferrin receptor-1 (*TFRC*) and divalent metal transporter-1 (solute carrier family 11 member 2 [*SLC11A2*]); storage, ferritin light chain (*FTL*) and ferritin heavy chain (*FTH1*); and export, ferroportin (*SLC40A1*); as well as the iron regulatory proteins, IRP1 (aconitase-1 [*ACO1*]) and IRP2 (iron responsive element binding protein-2 [*IREB2*]); and regulator of iron absorption/export, hepcidin (hepcidin antimicrobial peptide [*HAMP*]) at 24hpi in BCi-Ns1.1 cells and 5dpi in murine whole lung tissue. *TFRC* expression decreases in both BCi-Ns1.1 cells and murine lung (**Figure 1A,I**), and *SLC11A2* expression decreases in murine lung (**Figure 1B,J**), during IAV infection compared to sham-infected controls. *FTL* and *FTH1* expression decreases in BCi-Ns1.1 cells (**Figure 1C,D**) but increases in murine lung (**Figure 1L,M**) during infection. *SLC40A1* expression increases in both BCi-Ns1.1 cells (**Figure 1E**) and murine lung (**Figure 1M**) during IAV infection. *ACO1* and *IREB2* expression decreases, and *HAMP* expression increases, in BCi-Ns1.1 cells (**Figure 1F,H**), while these factors are not significantly altered in murine lung tissue (**Figure 1N,P**), during infection. We also show that IAV infection results in the increased number of cells with intermediate and high levels of intracellular iron accumulation in whole lung tissue during infection *in vivo* (**Figure S3A-D)**.

**Figure 1:**
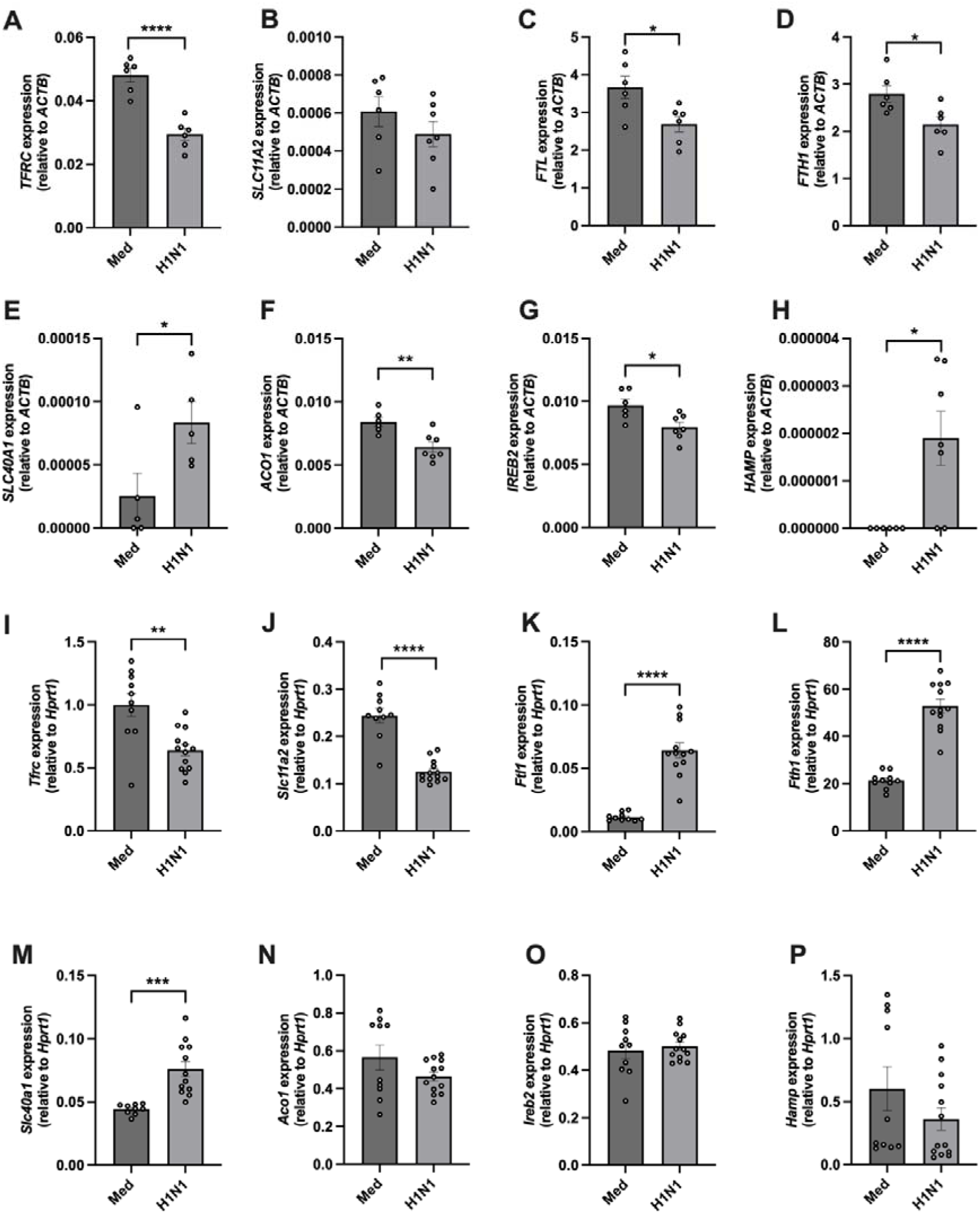
Influenza A virus infection alters iron metabolism in human AECs and murine whole lung tissue. BCi-Ns1.1 cells were infected with H1N1 (A/Auckland/1/2009) or sham-infected with media (Med). At 24 hours post-infection, gene expression of **A)** transferrin receptor-1 (*TFRC*), **B)** divalent metal transporter-1/solute carrier family 11 member 2 (*SLC11A2*), **C)** ferritin light chain (*FTL*), **D)** ferritin heavy chain (*FTH1*), **E)** ferroportin/solute carrier family 40 member 1 (*SLC40A1*), **F)** iron regulatory protein-1/aconitase-1 (*ACO1*), **G)** iron regulatory protein-2/iron responsive binding element protein-2 *(IREB2*) and **H)** hepcidin (*HAMP*) assessed in cells by qPCR. Mice were intranasally inoculated with H1N1 (A/Puerto Rico/8/1934) or sham-inoculated with media (Med). At 5 days post-infection, gene expression for **I)** *Tfrc*, **J)** *Slc11a2*, **K)** *Ftl1*, **L)** *Fth1*, **M)** *Slc40a1*, **N)** *Aco1*, **O)** *Ireb2* and **P)** *Hamp* was assessed in whole lung tissue by qPCR. Data (n=5-13) are presented as mean ± SEM. \**p*<0.05, \*\**p*<0.01, \*\*\**p*<0.001, and \*\*\*\**p*<0.0001.

Bulk RNA sequencing on fully differentiated primary human nasal AECs infected with different seasonal and emerging strains of influenza A and B viruses showed similar changes in expression for iron-related factors, with *TFRC*, *FTL* and *ACO1* expression consistently reduced with different infections as observed with H1N1 infection in BCi-Ns1.1 AECs (**Figure 2A-F**). Together these data show that human AECs and murine lung cells respond to different strains of influenza infection by significantly altering cellular and tissue iron metabolism.

**Figure 2:**
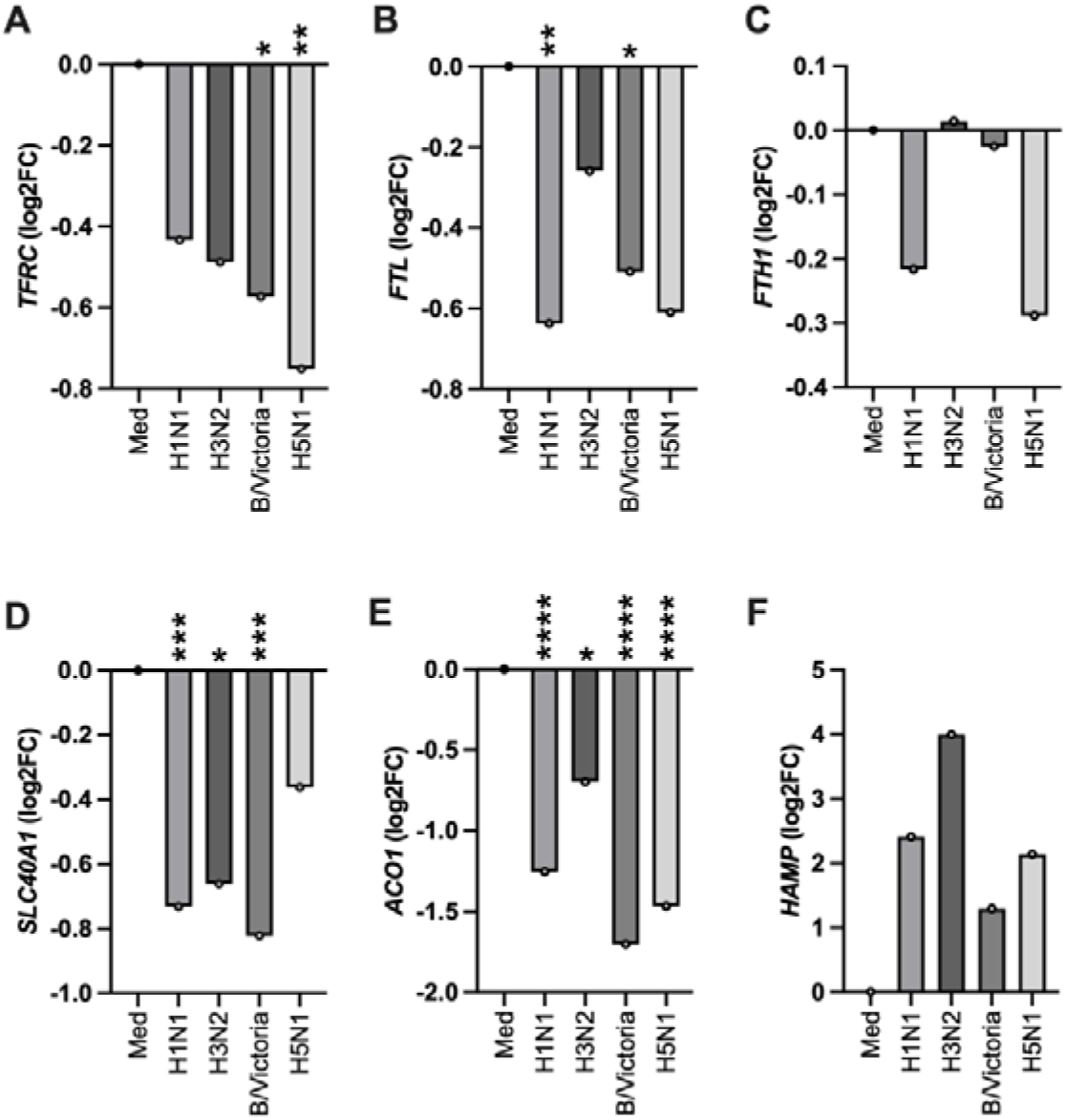
Seasonal and emerging strains of influenza viruses alter iron metabolism in human nasal airway epithelial cells (AEC). Nasal AECs were infected with H1N1 (A/Singapore/G2-25.1/2014), H3N2 (A/Singapore/CDC-204/2014), B/Victoria (B/Singapore/G2-14.1/2014), or H5N1 (A/Indonesia/CDC-1031/2007), or sham-infected with media (Med). At 24 hours post-infection **A)** transferrin receptor-1 (*TFRC*), **B)** ferritin light chain (*FTL*), **C)** ferritin heavy chain (*FTH1*), **D)** ferroportin/solute carrier family 40 member 1 (*SLC40A1*), **E)** iron regulatory protein-1/aconitase-1 (*ACO1*), and **F)** hepcidin (*HAMP*) gene expression was assessed in cells using RNA sequencing. Data (n=8) are presented as mean log2 fold change (FC) relative to sham-infected (Med) controls. Significance is presented where the FDR value is \**p*<0.05, \*\**p*<0.01, \*\*\**p*<0.001, and \*\*\*\**p*<0.0001.

### Modulating iron availability alters susceptibility to IAV infection in human AECs

Given that we show altered iron metabolic gene responses in AECs and lungs during influenza infections, we next sought to determine how altered iron availability may affect susceptibility to IAV infection. BCi-Ns1.1 cells were cultured with increasing concentrations of FAC (Fe^3+^; 10, 50 or 100μM) for 24 hours prior to infection with H1N1. FAC increases IAV replication in AECs in dose-dependent manner (**Figure 3A**; 24hpi). FAC-mediated increases in viral replication are observed as early as 12hpi (**Figure 3B**; 50μM FAC). To determine if these increases in IAV replication are dependent on the form of iron available and could be reversed, we next assessed the effects of treatment with holo-Tf or the iron chelator, DFO, alone or in combination with FAC, prior to IAV-infection. Holo-Tf (**Figure 3C**) increases IAV titer at 24hpi. Furthermore, DFO protects against IAV infection, reducing viral titer when given alone and preventing FAC-induced increases in IAV replication when given in combination (**Figure 3D**). Therefore, modulating iron availability has profound effects on IAV infectivity in AECs with increased ferric and Tf-bound iron increasing the susceptibility of AECs to infection. The profound effects of iron availability on infection did not correspond with any obvious effects on gene expression of innate host immune factors (**Figure S4**).

**Figure 3:**
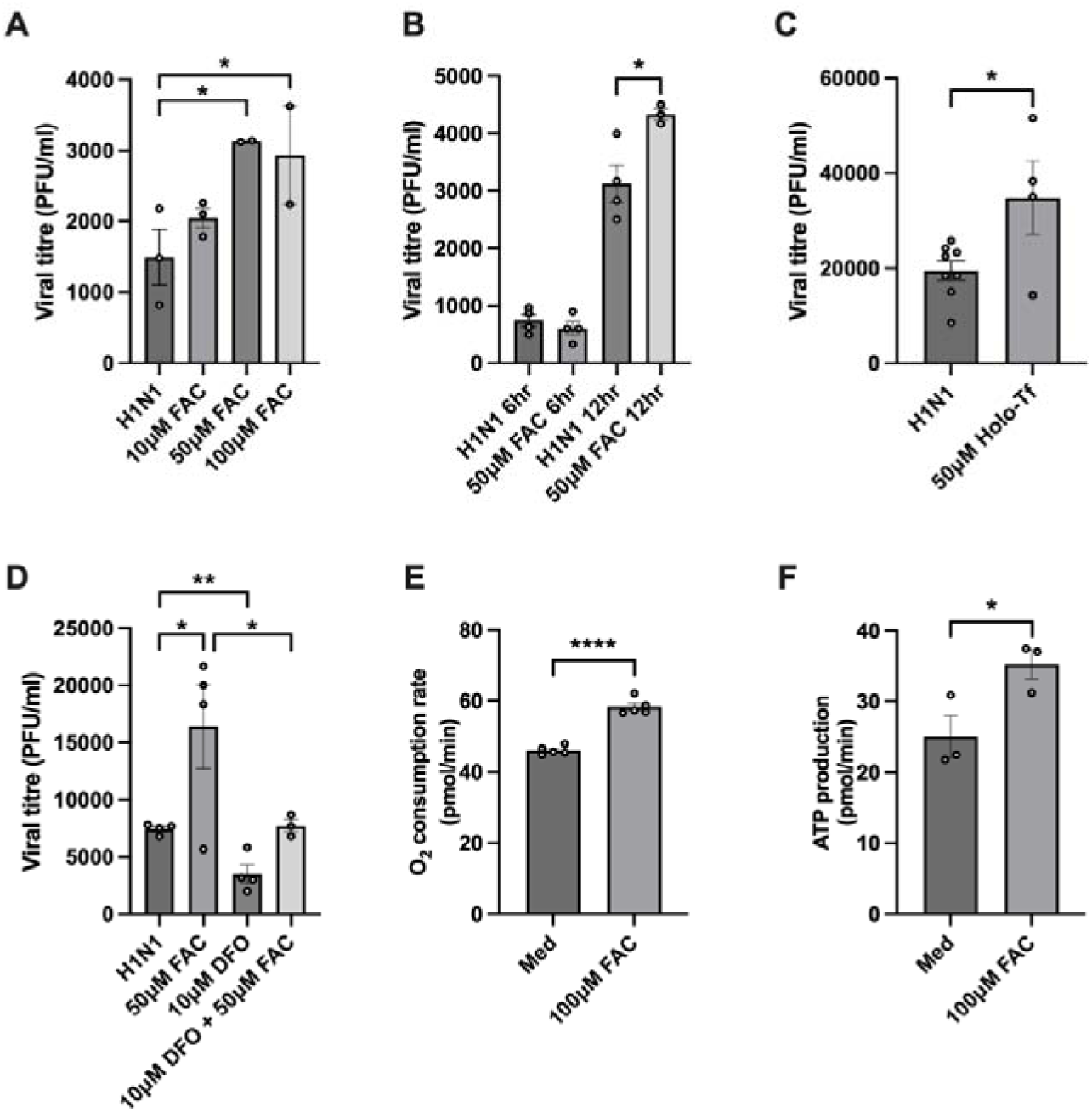
Modulating iron availability alters susceptibility to influenza A virus infection and cellular metabolism of human airway epithelial cells (AEC). BCi-Ns1.1 cells were cultured with iron modulating agents (10, 50, or 100μM) for 24 hours prior to infection with H1N1 (A/Auckland/1/2009) or sham-infected with media (Med). Viral titers in the supernatant were measured via plaque assay following exposure to: **A)** ferric ammonium citrate (FAC) at 24 hours post-infection (hpi) and **B)** 6 and 12hpi; **C)** holo-transferrin (Holo-Tf; 24hpi); **D)** deferoxamine (DFO; 24hpi). Primary human bronchial AECs were cultured with FAC (100μM) overnight prior to assessment of **E)** oxygen consumption rate and **F)** adenosine triphosphate production by Seahorse Cell Mito Stress assay. Data (n=2-15) are presented as mean ± SEM. \**p*<0.05, \*\**p*<0.01, \*\*\**p*<0.001, and \*\*\*\**p*<0.0001.

Interestingly, in separate experiments, we show that following 24hour culture with FAC (100 μM), oxygen consumption rate (OCR) and adenosine triphosphate (ATP) production in AECs is significantly increased (**Figure 3E,F**). These findings demonstrate that increased iron availability increases cellular metabolic activity in AECs, which may provide the energy required for increased viral replication.

### Increased iron loading and availability increases IAV infection and infection-induced disease *in vivo*

To assess the effect of increased iron levels on IAV infection and infection-induced disease outcomes *in vivo*, mice were fed a high iron diet (HID) for eight weeks (21), prior to infection with H1N1 (**Figure S2B**). Iron levels are increased in both the liver and, importantly, lungs of mice fed a HID. The total levels of non-heme iron are not greatly impacted in the lungs by IAV infection at 3dpi (**Figure 4A,B**), despite the increased intensity of iron loading observed in individual cells in IAV infection (**Figure S3**). IAV infection differentially effects iron-related genes in the liver compared to the lung (**Figure 4**, and described in supplement).

**Figure 4:**
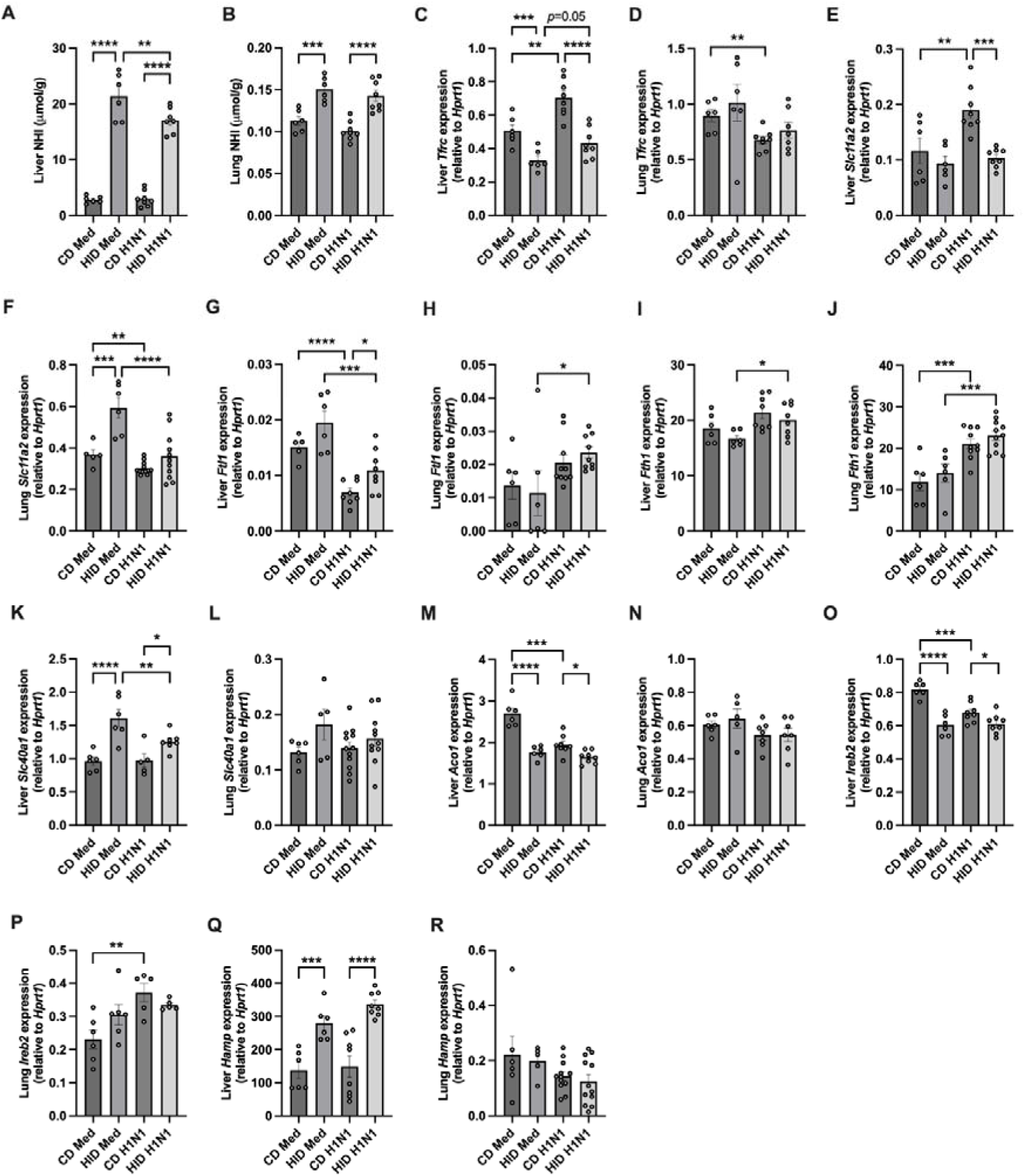
Influenza A virus (IAV) infection and high iron diet (HID)-induced increased iron loading alter iron metabolism systemically and in the lungs. Mice were fed a HID or control diet (CD) *ad libitum* for eight weeks prior to intranasal inoculation with H1N1 (A/Puerto Rico/8/1934) or sham-inoculation with media (Med). At 3 days post-infection, non-heme iron (NHI) levels were measured in **A)** liver and **B)** lung tissue. Gene expression was assessed in liver (**C, E, G, I, K, M, O, Q**) and lung (**D, F, H, J, L, N, P, R**) tissue by qPCR for: (**C-D)** transferrin receptor-1 (*Tfrc*); (**E-F**) divalent metal transporter-1/solute carrier family 11 member 2 (*Slc11a2)*; (**G-H**) ferritin light chain (*Ftl1)*; (**I-J**) ferritin heavy chain (*Fth1*); (**K-L)** ferroportin/solute carrier family 40 member 1 (*Slc40a1*); (**M-N**) iron regulatory protein-1/aconitase-1 (*Aco1*); (**O-P**) iron regulatory protein-2/iron responsive binding element protein-2 (*Ireb2*); and (**Q-R**) hepcidin (*Hamp*). Data (n=5-12) are presented as mean ± SEM. \**p*<0.05, \*\**p*<0.01, \*\*\**p*<0.001, and \*\*\*\**p*<0.0001.

Interestingly, we show that the total number and/or proportion (% of parent) of TFR1^hi^ alveolar and/or interstitial macrophages, monocytes, neutrophils and eosinophils is increased in whole lung tissues during IAV infection (**Figure 5**). TFR1^hi^ populations of these cells, as well as TFR1^hi^ epithelial and endothelial cells, are increased in during infection mice with increased HID-induced iron loading compared to IAV-infected controls (**Figure 5**).

**Figure 5:**
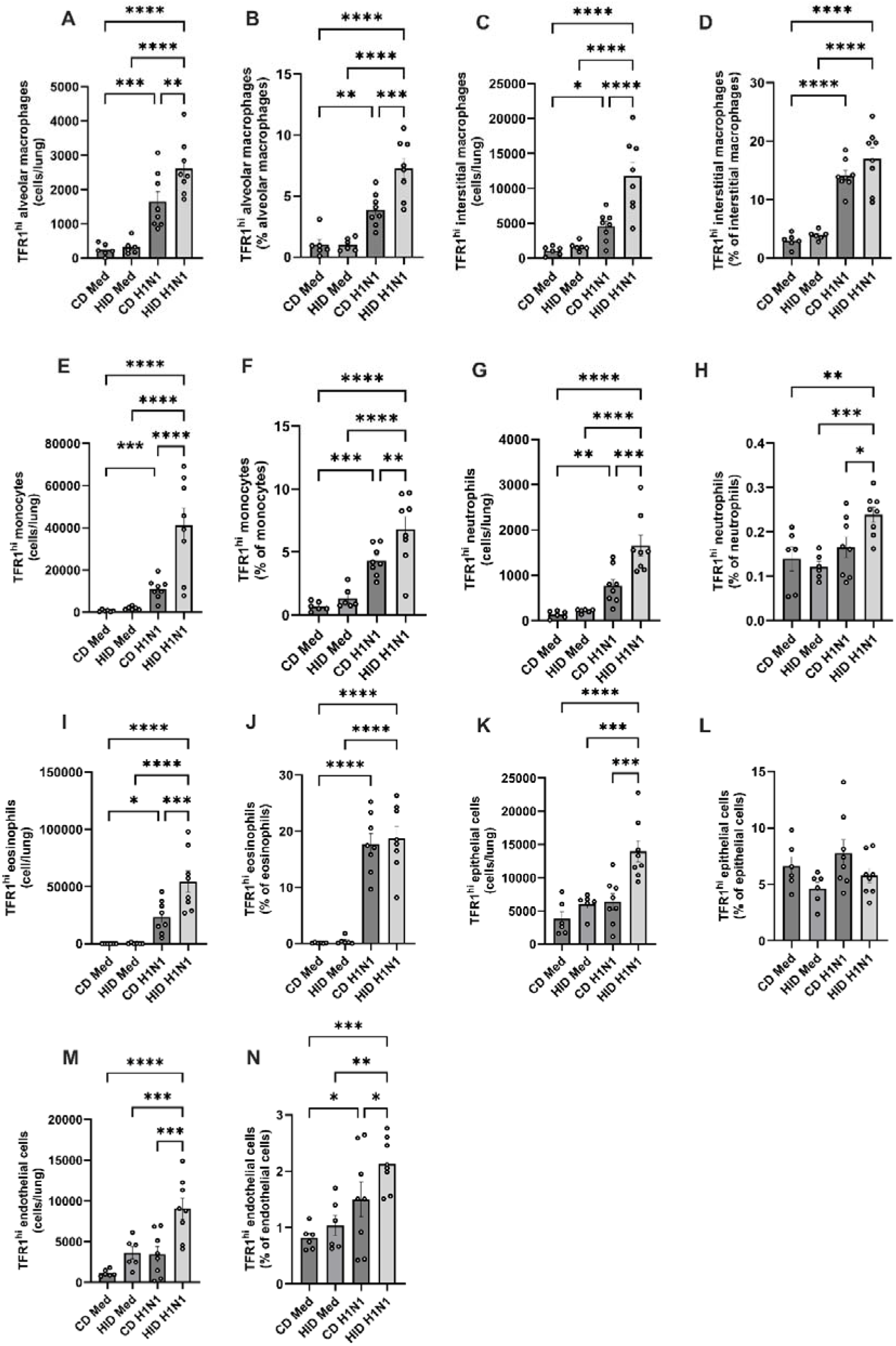
Increased iron loading increases transferrin receptor (TFR)1^hi^ cell numbers in the lungs in influenza A virus (IAV) infection. Mice were fed a high iron diet (HID) or control diet (CD) *ad libitum* for eight weeks prior to intranasal inoculation with H1N1 (A/Puerto Rico/8/1934) or sham-inoculation with media (Med). At 3 days post-infection (dpi), the total number and proportion (% of parent cell population) of TFR1^hi^ **A, B)** alveolar macrophages, **C, D)** interstitial macrophages, **E, F)** monocytes, **G, H)** neutrophils, **I, J)** eosinophils and **K, L)** epithelial and **M, N)** endothelial cells were assessed in whole lung tissue homogenates using flow cytometry. Data (n=6-8) are presented as mean ± SEM. \**p*<0.05, \*\**p*<0.01, \*\*\**p*<0.001, and \*\*\*\**p*<0.0001.

Importantly, HID-induced iron loading increases IAV-induced weight loss (**Figure 6A**), airways resistance, both at baseline (**Figure 6B)** and in response to methacholine challenge (**Figure 6C),** viral titer (**Figure 6D**), and total inflammatory cell and neutrophil numbers in bronchoalveolar lavage fluid (BALF, **Figure 6E, F**), at 5dpi. HID-induced increases in airways resistance and inflammatory cell numbers in BALF were also seen at 3dpi (**Figure S5**).

**Figure 6:**
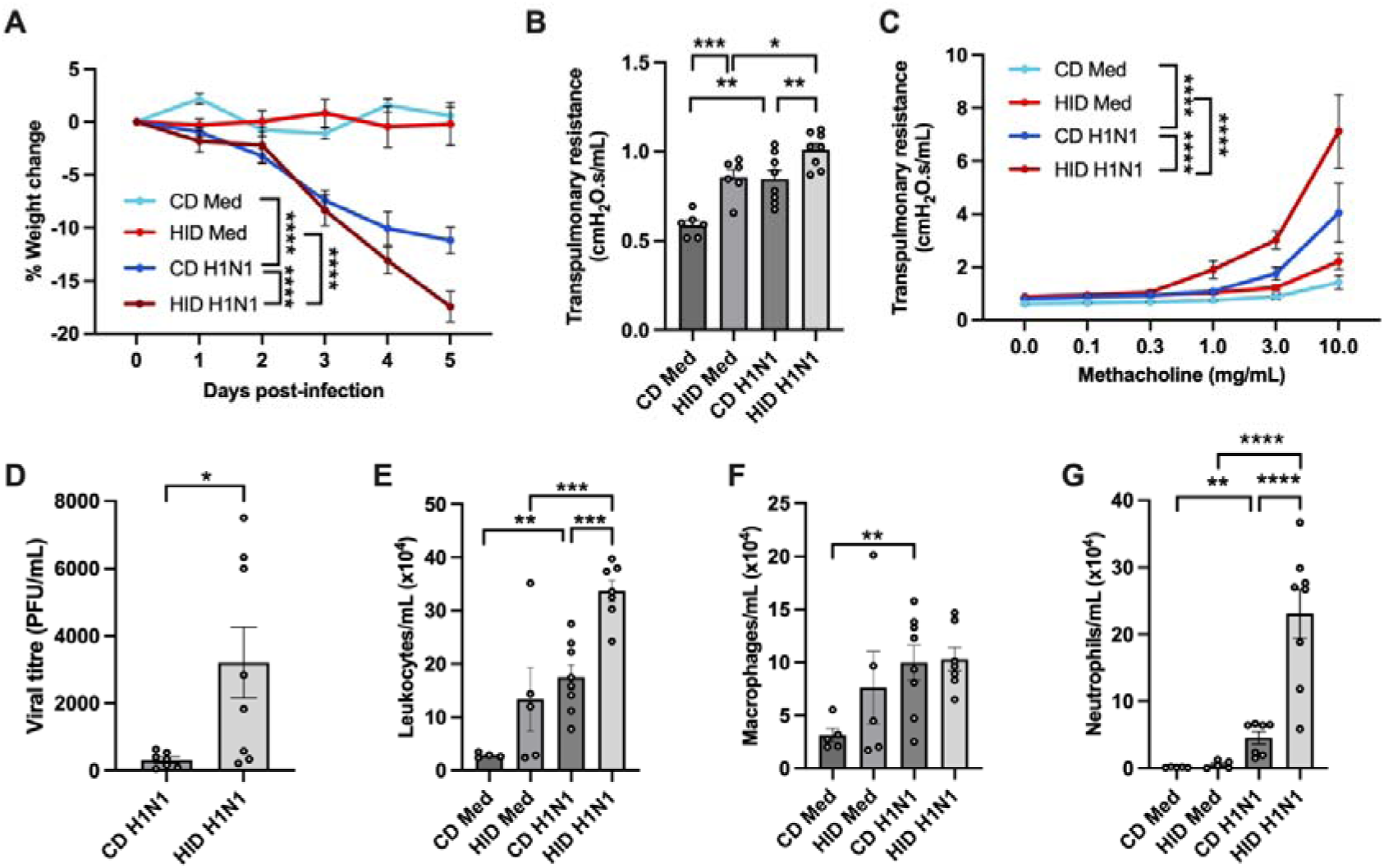
Increased iron loading increases influenza A virus (IAV) infection and infection-induced disease. Mice were fed a high iron diet (HID) or control diet (CD) *ad libitum* for eight weeks prior to intranasal inoculation with H1N1 (A/Puerto Rico/8/1934) or sham-inoculation with media (Med). At 5 days post-infection (dpi), mice were assessed for **A)** % weight change from 0dpi, lung function in terms of transpulmonary resistance at **B)** baseline and **C)** during challenge with increasing doses of nebulized methacholine, **D)** viral titer in bronchoalveolar lavage fluid (BALF; in plaque forming units [PFU]) and **E)** total leukocytes, **F)** macrophages and **G)** neutrophils in BALF. Data (n=6-8) are presented as mean ± SEM. \**p*<0.05, \*\**p*<0.01, \*\*\**p*<0.001, and \*\*\*\**p*<0.0001. Statistical significance is shown at 5dpi for % weight change (**A**) and 10mg/mL methacholine for airways hyperresponsiveness (**C**).

Together these findings demonstrate that IAV infection and iron loading exert potent effects on iron metabolism both systemically and in the lungs, and increased systemic and lung iron loading significantly worsens IAV infection and infection-induced disease during early. More severe disease observed during infection with increased iron loading is associated with increases in the number and proportion of TFR1^hi^ cell populations in the lungs.

### Inhibition of TFR1 responses protects against IAV infection and infection-induced disease *in vivo*

We show that IAV titer increases in response to increased concentrations of holo-Tf and that *TFRC* expression is reduced, but TFR1^hi^ populations of cells is increased, in response to IAV infection. TFR1^hi^ populations are even further increased in mice with increased iron loading that have more severe disease. These findings suggest that TFR1-mediated uptake of Tf-bound iron may increase IAV infection and that altered TFR1 responses play an important role in mediating iron-associated host susceptibility to IAV infection and disease.

To investigate the role of lung TFR1 responses in IAV infection and infection-induced disease, mice were intranasally administered anti-TFR1 antibody (αTFR1) to deplete TFR1-mediated responses during H1N1 infection (**Figure S1C**). Disease outcomes were assessed at 5dpi (**Figure S1C**). Administration of αTFR1 reduces IAV-induced weight loss (**Figure 7A**), airways resistance, both at baseline (**Figure 7B)** and in response to methacholine challenge (**Figure 7C)**, viral titer (**Figure 7D**), and total leukocyte, macrophage and neutrophil numbers in the airways (**Figure 7E-F**), at 5dpi. αTFR1-mediated decreases in inflammation during infection are also observed at 3dpi (**Figure S6**).

**Figure 7:**
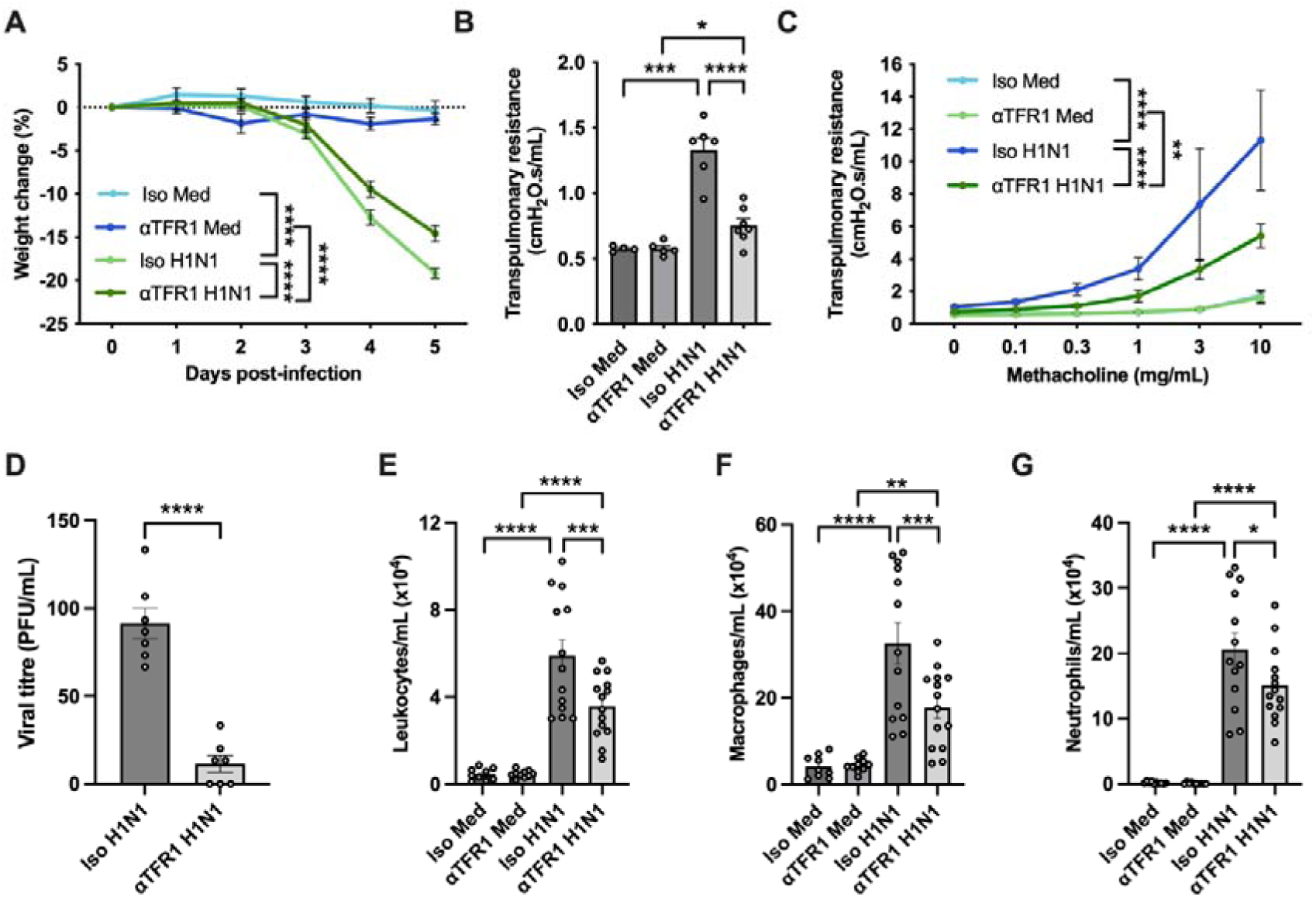
Inhibition of TFR1 responses protects against influenza A virus infection and infection-induced disease. Mice were intranasally inoculated with H1N1 (A/Puerto Rico/8/1934) or sham-inoculated with media (Med) and intranasally administered anti-transferrin receptor-1 (aTFR1; 1mg/kg) or isotype control (Iso; 1mg/kg) monoclonal antibody on days 0, 2 and 4. At 5 days post-infection (dpi), mice were assessed for **A)** % weight change from 0dpi, lung function in terms of transpulmonary resistance at **B)** baseline and **C)** during challenge with increasing doses of nebulized methacholine, **D)** viral titer in bronchoalveolar lavage fluid (BALF; in plaque forming units [PFU]), and **E)** total leukocytes, **F)** macrophages and **G)** neutrophils in BALF. Data (n=4-14) are presented as mean ± SEM. \**p*<0.05, \*\**p*<0.01, \*\*\**p*<0.001, and \*\*\*\**p*<0.0001. Statistical significance is shown at 5dpi for % weight change (**A**) and 10mg/mL methacholine for airways hyperresponsiveness (**C**).

### Increased iron loading increases, whilst inhibition of TFR1 responses decreases, the expression of specific inflammasome-related genes during IAV infection *in vivo*

We next sought to understand how iron-modifying interventions may be affecting disease outcomes from an immune perspective by measuring the expression of inflammasome-related factors in lung tissue during iron loading or in response to αTFR1 administration at 3 and 5dpi. IAV infection increased the expression of *Aim2*, *Nlrc4*, *Nlrp3*, *Casp1*, *Casp4*, *Il1b*, and *Il18*, at both 3 (**Figure 8**) and 5dpi (**Figure S7**). At 3dpi, HID-induced iron loading increased *Aim2* expression during infection, compared to mice fed a CD (**Figure 8A**). αTFR1 administration reduced *Aim2* expression during infection, compared to mice treated with isotype control antibody (**Figure 8B**). Similarly, *Nlrc4* expression was increased by iron loading and reduced by αTFR1 administration during infection (**Figure 8C,D**). *Nlrp3* expression was not altered by iron loading or αTFR1 administration at 3dpi (**Figure 8E,F**), however, iron loading significantly reduced *Nlrp3* expression at 5dpi (**Figure S7**). *Casp1* expression was not significantly altered by iron loading but was reduced by αTFR1 administration during infection (**Figure 8G-H**). Iron loading increased *Casp4* expression during infection at 3dpi (**Figure 8I**) but αTFR1 administration had no affect (**Figure 8J**). *Il1b* expression was not significantly altered by iron loading or αTFR1 administration (**Figure 8K, L**). *Il18* expression was increased by iron loading (**Figure 8M**) and reduced by αTFR1 administration at 3dpi (**Figure 8N**). Most of the iron overload-or αTFR1-induced changes in inflammasome gene expression were not observed at 5dpi (**Figure S7**).

**Figure 8:**
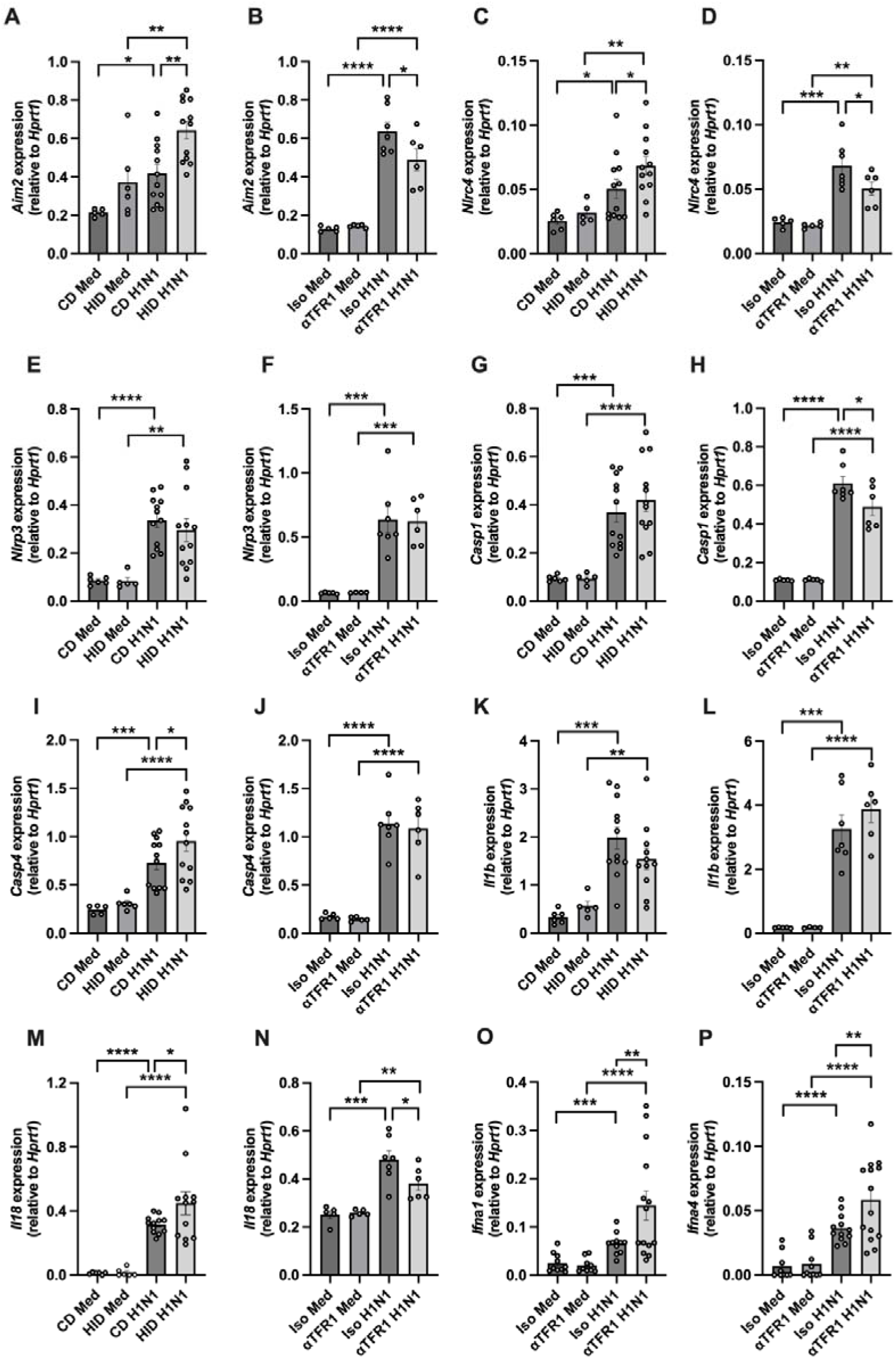
Increased iron loading or anti-transferrin receptor (TFR)1 mononclonal antibody (αTFR1) treatment affects the expression of inflammasome and type I interferon (IFN) responses during influenza A virus infection. Mice were fed a high iron diet (HID) or control diet (CD) *ad libitum* for eight weeks prior to intranasal infection with H1N1 (A/Puerto Rico/8/1934) or sham-infected with media (Med). A separate group of mice were intranasally infected with H1N1 or sham-infected (d0) and intranasally administered anti-transferrin receptor-1 (αTFR1; 1mg/kg) or isotype control (Iso; 1mg/kg) monoclonal antibody on days 0 and 2. At 3 days post-infection (dpi), gene expression was assessed in lung tissue by qPCR for **A, B)** absent in melanoma-2 (*Aim2*), **C, D)** NLR family CARD domain containing 4 (*Nlrc4*), **E, F)** NLR family pyrin domain containing 3 (*Nrlp3*), **G, H)** caspase-1 (*Casp1*), **I, J)** caspase-4 (*Casp4*), **K, L)** interleukin-1 beta (*Il1b)*, and **M, N)** interleukin-18 (*Il18*). At 5dpi, gene expression for **O)** interferon alpha 1 (*infa1*) and **P)** interferon alpha 4 (*infa4*) were also assessed in IAV-infected and sham-infected mice, treated with αTFR1 or Iso. Data (n=5-12) are presented as mean ± SEM. \**p*<0.05, \*\**p*<0.01, \*\*\**p*<0.001, and \*\*\*\**p*<0.0001.

### Inhibition of TFR1 increases type I IFN responses during IAV-infection *in vivo*

We next measured the expression of key anti-viral factors involved in IAV clearance. IAV infection increased the expression of *Ifnb1*, *Ifna1*, and *Ifna4* at both 3 (**Figure S8)** and 5dpi (**Figure 8O,P** and **Figure S9**). αTFR1 administration, significantly increases *Ifna1* and *Ifna4* expression during IAV infection at 5dpi (**Figure 8O,P**). HID-induced iron loading significantly increases *Ifna4* expression during IAV infection at 5dpi (**Figure S9**). The effects of αTFR1 on *Ifna1* and *Ifna4* expression during IAV infection was not observed at 3dpi (**Figure S8**).

## DISCUSSION

Using complementary *in vitro* and *in vivo* studies, we demonstrate how IAV infection alters iron metabolism in the lung and systemically during the early stages of infection, and that altering iron levels and metabolism in the cells and tissues of the respiratory tract and/or systemically profoundly affects susceptibility to infections and resultant IAV-induced disease.

Previous studies have reported associations between dysregulated iron homeostasis and the severity of respiratory viral infections, including SARS-CoV-2 and IAV. High serum ferritin is associated with severe respiratory failure in COVID-19 and has been identified as a predictive biomarker for the triage and management of this disease (10–12). Low serum iron and high ferritin levels also correlate with disease severity during H7N9 infection (31). Several viruses, including IAV and SARS-CoV-2, have been recently shown to exploit TFR1 for viral entry (32–34). However, few studies have investigated the functional nature of the interplay between altered iron metabolism and host responses to viral infection and how this interplay affects IAV infection and infection-induced disease outcomes.

Here we have conducted a comprehensive series of analyses to show that IAV infections decrease the expression of genes for factors involved in cellular iron uptake, TFR1 and divalent metal transporter-1/SLC11A2, both in lung tissue *in vivo* and in human AECs. IAV infections also increase the expression of cellular iron exporter, ferroportin/SLC40A1, both in the lung *in vivo* and in human AECs. We also show that IAV infection increases the expression of TFR1 and divalent metal transporter-1/SLC11A2 in the liver, demonstrating that the host responds to IAV infection by differentially modulating iron metabolism in the lungs *versus* systemically. Importantly, IAV infection results in a significant increase in iron accumulation in cells in lung tissues, which corresponds with an increased in ferritin gene expression in whole lung tissue. Furthermore, we show that infection increases the number and proportion of TFR1^hi^ immune, epithelial and endothelial cells *in vivo.* Together, these findings demonstrate that modulation of cellular iron metabolism is a significant feature of the host-pathogen response to IAV.

We extend upon our findings to show that increased iron availability, both in AECs and *in vivo*, increases susceptibility to IAV infection. Importantly, iron-chelation protects against IAV infection. Interestingly, we show that increased iron levels in AECs have little effect on the expression of innate immune factors (**Figure S3**), but do have a potent effect on cellular metabolism, suggesting that increased iron availability may enhance cellular metabolism required to facilitate IAV replication in AECs as opposed to affecting AEC antiviral immunity. We also show increased iron loading *in vivo* is associated with more severe infection-induced pathology, with mice fed a HID, which increases systemic and lung levels of iron, increasing weight loss, airflow obstruction and airways inflammation during infection. Together, these findings demonstrate that increased iron availability has profound effects on IAV infection and infection-induced outcomes during the early stage of infection.

Increased disease severity in mice with increased iron loading is associated with significant increases in the number and proportion of TFR1^hi^ cells in the lungs during IAV infection. Notably, we show that local administration of αTFR1 during infection reduces IAV titer, weight loss and inflammation and improves lung function *in vivo*. Given that others have shown that IAV is capable of exploiting TFR1 for entry into host cells (32) and we have shown that increasing iron availability, with ferric or TF-bound iron, increases IAV replication in AECs, pharmaceutical neutralization of TFR1 may protect against IAV infection and infection-induced disease by both inhibiting TFR1-mediated viral entry and reducing intracellular iron availability. αTFR1 may also mediated protective responses through suppression of TFR1^hi^ immune cells, which are induced by IAV infection and are increased in more severe disease, given that we and others show that TFR1^hi^ immune cells have altered immune phenotypes (21, 23, 35, 36). Together these findings demonstrate that alterations in TFR1 play a crucial role in mediating host defence against IAV infection and this likely occurs through different mechanisms in different cell populations in the lungs and airways.

Interestingly, our data show that inflammasome and type I IFN responses may be affected by altered iron metabolism during IAV infection in whole lung tissues, which contain both immune and mesenchymal cells. Iron loading increases, while αTFR1 administration decreases, *Aim2*, *Nlrc4*, caspase and *Il18* expression in the lung during infection. Inflammasome responses are critical for the pathogenesis of IAV infections (37–39). Previous studies have shown that AIM2 deficiency attenuates lung injury and improves survival following IAV infection *in vivo* (37). Interestingly, NLRC4 activity has been shown to regulate dendritic cell phenotypes in the lung, influencing the magnitude of T-cell responses during IAV infection (40), however, how IAV infections activate NLRC4 has yet to be determined. Additionally, caspase-4 has been shown to contribute to the induction of pyroptosis during IAV infection (41, 42). We also show that αTFR1 treatment boosts type I IFN responses in the lung during infection, which play key roles in protecting against the early stages of IAV infection (43, 44). Together these findings highlight potentially important roles for iron metabolism in modifying inflammasome responses mediated by AIM2, NLRC4, caspase-1 and/or caspase-4, and anti-viral responses mediated by type I IFNs during IAV infection that may contribute to the development of disease.

Subjects with asthma are more susceptible to severe IAV infections (45–48) and we have previously shown that subjects with asthma have increased iron accumulation and TFR1 responses in their AECs and lung tissues (2, 21, 49). Given that we show that increased iron and TFR1 responses in the airways increases susceptibility to IAV infection and are associated with more severe disease, we propose that altered iron homeostasis in the airways of subjects with asthma may play an important role in increased susceptibility to IAV infection. Furthermore, given that COPD is also associated with increased iron accumulation in the airways (50–54), we also propose a similar role for altered iron metabolism in increasing susceptibility to IAV in COPD. Importantly, therapeutic iron chelation or neutralisation of TFR1 in the airways may be particularly effective treatments for IAV infections in subjects with asthma and COPD. A recent study identified correlations between iron redistribution and persisting symptomatology of SARS-CoV-2 infections, with increased monocyte iron loading, low serum iron, anaemia, lymphopenia, and increased iron demand of lymphocytes seen in individuals with long COVID (55). This observation suggests that treatments that target iron metabolism and cellular distribution may also be effective in alleviating the symptoms of multiple viral infections, particularly chronic SARS-CoV-2.

IAV respiratory infections remain a major threat to human health. Circulating viruses rapidly mutate and new viruses emerge, rendering vaccines and antiviral treatments obsolete. Strategies that boost protective, or reduce pathogenic, host responses may provide broadly effective, long-term therapeutic options. We have identified a key role for iron status in modifying host responses to IAV that can be harnessed to protect against IAV infection and associated disease during the early stages. Further research is required to understand whether iron targeting therapies may be broadly applicable, not only to other strains of IAV, such as emerging strains and variants of concern, but also to other respiratory viral infections. Importantly, whether iron-targeting therapies can be used for protecting against severe viral infections in subjects with chronic respiratory diseases such as asthma and COPD requires further investigation.

## ACKNOWLEDGEMENTS

Generation of BCi-Ns1.1 data was supported by grant funding awarded by the Hunter Medical Research Institute, Australia, to JRM. The work for the generation of human nasal AEC RNA sequencing data was supported by grants awarded by the National Medical Research Council, Singapore, to DYW (NMRC/CIRG/1458/2016) and to KST (MOH-OFYIRG19may-0007). ALP was supported by an Australian Government Research Training Program Scholarship. The authors would like to thank undergraduate students, Mia J. Gottstein, Lily M. Cassano and Kane Predebon (University of Newcastle), for their contributions to the investigation and analysis, conducted under the supervision of JRM and JCH, Tegan Hunter (research assistant, University of Newcastle) for her technical support, and the staff of the University of Newcastle BioResearch Facilities for their operational support.

## Notes

### Competing Interest Statement

The authors have declared no competing interest.

