## Supplemental methods and data for "Crucial role for iron metabolism in mediating influenza A virus infection and associated disease"

**ORIGINAL RESEARCH ARTICLE**

### SUPPLEMENTARY METHODS

#### Viruses

A/Auckland/1/2009 (H1N1) and A/Puerto Rico/8/1934 (H1N1) were sourced from the WHO Collaborating Centre for Reference and Research on Influenza (Parkville, Victoria, Australia). A/Singapore/G2-25.1/2014 (H1N1), A/Singapore/CDC-204/2014 (H3N2), B/Singapore/G2-14.1/2014 (Victoria lineage), and A/Indonesia/CDC-1031/2007 (H5N1) were clinical isolates obtained from infected patients.

#### IAV infections in human airway epithelial cells (AEC)

Minimally immortalised bronchial AECs (clonal population of basal cell immortalized nonsmoker 1 cell line; BCi-Ns1.1) were obtained from Prof. Ronald Crystal's lab at Weill Cornell Medicine and Cornell University (NY, USA). BCi-Ns1.1 cells were cultured in BEBM™ Bronchial Epithelial Cell Basal Medium (BEBM) supplemented with BEGM™ Bronchial Epithelial Cell Growth Medium SingleQuots™ Supplements and Growth Factors (Lonza, Australia), Amphotericin B (2.5µg/mL; Sigma-Aldrich, Germany), and penicillin:streptomycin (50units/mL:50µg/mL; Gibco, Australia; complete BEGM; 37˚C, 5% $CO_2$). Cells were detached with 0.05% Trypsin-EDTA (Gibco) then seeded into 24 well tissue culture-treated plates (CORNING, Australia) at $1x10^5$ cells/well in complete BEGM (1mL) and incubated for 24 hours (37˚C, 5% $CO_2$). Media were replaced with BEBM supplemented with BEGM™ Bronchial Epithelial Cell Growth Medium SingleQuots™ Supplements and Growth Factors, excluding hydrocortisone (Lonza); insulin, transferrin and selenium (ITS; 10µg/mL recombinant human insulin, 5.5µg/mL human transferrin, 5ng/mL sodium selenite; Sigma-Aldrich); linoleic acid (4.25µg/mL; linoleic acid-albumin from bovine serum albumin; Sigma-Aldrich); penicillin:streptomycin (50µg/mL; Gibco); and amphotericin B (2.5µg/mL; Sigma-Aldrich) to make BEBM minimal media, with or without ferric ammonium citrate (FAC; 10µM

[2.62μg/mL], 50μM [13.1μg/mL], 100μM [26.2μg/mL], Sigma-Aldrich), holo-transferrin (holo-Tf; 50μM; Sigma-Aldrich or deferoxamine (DFO; 10μM; Sigma-Aldrich) and cells incubated for 24 hours (37˚C, 5% $CO_2$). Media were then removed, and cells infected with H1N1 (A/Auckland/1/2009; multiplicity of infection [MOI] 0.5) in BEBM minimal media (1mL). After 1 hour, viral inoculum was replaced with fresh BEBM minimal media (1mL), and culture continued (37˚C, 5% $CO_2$). Cells and supernatant were collected 2, 6, 12 or 24 hours post-infection (hpi). Supernatant was stored at -80˚C for plaque assay. Cells were collected in Qiazol® or RLT buffer (QIAGEN, Germany) for RNA extraction.

**IAV infections in mice**

Six to eight week-old, specific pathogen free (SPF), female, wild-type BALB/c mice (obtained from the Animal Services Unit of the University of Newcastle, Callaghan, NSW, Australia) were housed in individually ventilated cages under SPF, PC2 conditions and a 12 hour light-dark cycle in a temperature (22 ± 2°C) and humidity (30–70%) controlled room (Hunter Medical Research Institute, New Lambton Heights, NSW, Australia). Food and water were provided *ad libitum*. Mice were intranasally inoculated with IAV (A/Puerto Rico/8/1934 [H1N1]; 7.5PFU in 50µl ultra-MDCK medium) under isoflurane anaesthesia on day 0 (**Figure S1A**). Control mice were sham infected with ultra-MDCK medium (50µl). To model increased systemic iron loading, mice were fed a high iron diet (HID; 2% carbonyl iron [20g Fe/kg] diet; Specialty Feeds, WA, Australia) for 8 weeks prior to infection (**Figure S1B**). Control mice were fed a control diet (CD; Specialty Feeds) with the same base composition, excluding ferrous iron level. To deplete TFR1-mediated cellular iron uptake, mice were treated with anti-mouse transferrin receptor-1 antibody (aTFR1; 2.5mg/kg; clone R217.1.3/TIB-219; BioXcell, United States) intranasally on days 0, 2 and 4 (**Figure S1C**). Control mice were administered isotype control antibody (Iso; IgG2a; clone 2A3; BioXcell). Mice were weighed daily. Lung function was measured at 3 and 5 days post-infection (dpi). At the endpoint, mice were euthanised by sodium pentobarbitone (Lethabarb; Virbac, Australia) overdose and bronchoalveolar lavage fluid (BALF), lung and liver tissues were collected for further analyses.

**RNA isolation and reverse transcription**

Excised murine lung and liver tissue was snap frozen in liquid nitrogen and stored at -80°C for RNA isolation using TRIzol® (Thermo Fisher, Australia) reagent. Tissue was homogenised in TRIzol (1mL; Thermo Fisher). Samples were centrifuged (10 min, 12000 x g, 4°C) and supernatant combined with chloroform (250µl; Sigma-Aldrich). After briefly vortexing, samples were incubated at room temperature for 10 min, then centrifuged (15 min, 12,000 x g, 4°C). The aqueous phase was aspirated and combined with ice-cold isopropanol (1:1; Sigma-Aldrich). Samples were vortexed, incubated (10 min, room temperature) and centrifuged (10 min, 12000 x g, 3°C) and supernatant discarded. RNA pellets were washed three times with molecular grade ethanol (75%; 1mL; Sigma-Aldrich) and centrifuged (5 min, 8000 x g, 4°C). Residual ethanol was aspirated, and samples air-dried on ice for up to 15 min. Pellets were resuspended in nuclease-free water (UltraPure DNase/RNase-Free Distilled Water, Invitrogen, United States).

RNA was isolated from BCi-Ns1.1 cells using RNeasy Mini kits (QIAGEN, Germany) according to the manufacturer's instructions. RNA concentration was determined using the NanoDrop™ 1000 Spectrophotometer (Thermo Fisher). RNA from lung and liver tissue (1000ng; in 8µl) was reverse transcribed using MMLV™ Reverse Transcriptase enzyme and reagents as follows (reagents and protocol from Thermo Fisher). Samples were treated with DNAse I amplification grade (1µl) and 10X DNAse reaction buffer (1µl) and incubated at room temperature for 15 min. DNAse STOP solution (1µl) was added and samples incubated at 65°C for 10 min using a Bio-Rad T100™ thermal cycler. Random hexamer primers (50ng/µl; 1µl) and dNTPs were added (10mM; 1µl) and samples incubated at 65°C for 5 min. A mix of 5X

First-Strand Buffer (4µl), 1,4-dithiothreitol (DTT; 100mM; 2µl) and nuclease-free $H_2O$ (1µl) was then added, and samples incubated at 37°C for 2 min. MMLV™ RT enzyme (1µl) was added and samples incubated at 25°C for 10 mins, 37°C for 50 mins and 70°C for 15 mins. cDNA was diluted in nuclease-free $H_2O$ (100µl) and stored at -30°C. cDNA was synthesized from BCi-Ns1.1 cell RNA (200ng) using the High-Capacity cDNA Reverse Transcription Kit (Applied Biosystems, United States) as per the manufacturer's instructions.

**Real-time quantitative PCR (qPCR)**

mRNA expression was assessed by SYBR Green qPCR (Bio-Rad, United States) using custom designed primers (**Table S1 and S2**) or TaqMan qPCR Assays (murine *Ifnb1*: Mm00439552_s1; *Hprt1*: Mm00446968_m1; Applied Biosystems) using CFX96 and CFX384 Touch Real-Time PCR Detection Systems (Bio-Rad) as per the manufacturer's guidelines. Relative expression was calculated to *Hprt1* (murine targets) or *ACTB* (human targets) using the $2^{-\Delta\Delta Ct}$ method:

$$\text{Relative expression} = 2^{\wedge -\,[Ct\,(TARGET)] - [Ct\,(HOUSEKEEPER)]}$$

**Measurement of viral titer by plaque assay**

Plaque assay was performed on mouse BALF or BCi-NS1.1 cell supernatant. MDCK-SIAT cells (Sigma-Aldrich) were cultured in Dulbecco's modified eagle medium (DMEM), supplemented with foetal bovine serum (10%; CellSera, Australia), sodium pyruvate (1mM; Sigma-Aldrich), sodium bicarbonate (1.8mg/mL; Sigma-Aldrich), penicillin:streptomycin (100units/mL:100µg/mL; Gibco), L-Glutamine (2mM; Gibco), HEPES (20mM; Gibco) and Geneticin™ (1mg/mL; Gibco). MDCK-SIAT cells ($4.5x10^5$/well) were seeded into 6 well tissue culture-treated plates and cultured overnight (37˚C, 5% $CO_2$). Cell monolayers were washed with PBS (Sigma-Aldrich), inoculated with viral samples serially diluted in Leibovitz's

L-15 medium (300µl; Thermo Fisher) supplemented with HEPES (20mM) and trypsin-TPCK (2µg/mL; Sigma-Aldrich), and incubated for 1 hour (37°C, 5% $CO_2$). Viral inoculum was removed and a semi-solid agarose gel layer (3mL; Leibovitz's L-15 medium supplemented with agarose [0.9%; Sigma-Aldrich], bovine serum albumin [1%; Sigma-Aldrich], sodium pyruvate [1mM], sodium bicarbonate [2.1mg/mL], penicillin:streptomycin [100units/mL:100µg/mL], HEPES [20mM] and trypsin-TPCK [2µg/mL]) placed over the monolayer. Cells were incubated for 3-4 days (37°C, 5% $CO_2$) and viral titre calculated using the formula:

*(Plaque number x dilution factor of viral inoculum) / inoculum volume in mL*

**Measurement of cellular metabolism by Seahorse assay**

Oxygen consumption rate and ATP production in primary human bronchial epithelial cells (HBECs, LONZA) was determined using a Seahorse XFe96 Analyzer (Agilent, United States). Human bronchial AECs ($2 \times 10^4$ cells/well) were expanded in submerged cultures using bronchial epithelial basal media (BEBM) supplemented with BEGM SingleQuotsTM (LONZA) for 72 hours, then cultured overnight in Seahorse XF cell culture microplates, prior to the assay. On the day of the assay, media was replaced with BEBM supplemented with sodium pyruvate (1mM), L-glutamine (2mM) and glucose (10mM) (Gibco) at 37°C. A Seahorse Cell Mito Stress assay kit (Agilent) was used to measure basal respiration and ATP production after the addition of oligomycin (1 µM), carbonyl cyanide 4-(trifluoromethoxy) phenylhydrazone (FCCP; 0.5µM) and antimycin A and rotenone (Rot/AA; 0.5µM), as per manufacturer's instructions. At basal level and after the addition of each compound, measurements were taken. To assess the effect of iron on these cellular parameters, cells were treated with 100µM of ferric ammonium citrate (FAC; Sigma-Aldrich) overnight prior to the assay.

**Measurement of lung function in mice**

Mice were anaesthetised with ketamine (100mg/kg; Parnell) and xylazine (10mg/kg; Troy Laboratories, Australia; intraperitoneal injection) prior to tracheotomy. Mice were intubated and ventilated at 450 breaths/min and tidal volume of 8mL/kg. transpulmonary resistance (Rrs) was measured at baseline and in response to increasing doses of nebulised methacholine (Mch; 0, 0.1, 1, 3, 10mg/mL; 15μl) using the FlexiVent apparatus (FX Module 1 System; Scireq, Canada). Data was recorded using FlexiWare software v7.6.

**Measurement of inflammatory cell numbers in BALF**

The left lung lobe was lavaged with DMEM (2 x 500μl). Erythrocytes were removed by incubation in red blood cell lysis buffer (155mM $NH_4Cl$, 12mM $NaHCO_3$, 0.1mM ethylenediaminetetraacetic acid [EDTA], pH 7.35; 1mL; 5 min; on ice). Viable cells/mL were calculated by trypan blue exclusion using a haemocytometer. Cells were cytocentrifuged (15 x *g*, 10 min; Shandon Cytocentrifuge 2, Marshal Scientific, United States) onto by slides, air dried overnight and stained with May-Grunwald and Giemsa. Different cell populations were identified and enumerated based on cellular morphology and staining under light microscopy (40x magnification).

### SUPPLEMENTARY RESULTS

**IAV infection and increased iron loading have differential effects on iron-related gene expression and iron metabolism in the liver and lungs**

IAV infection increases liver *Tfrc* expression, and decreases lung *Tfrc* expression, in mice fed a CD (**Figure 4C,D**). HID-induced iron loading decreases liver *Tfrc* expression but does not affect lung *Tfrc* expression (**Figure 4C,D**). Similarly, IAV infection increases liver

*Slc11a2* expression, and decreases lung *Slc11a2* expression (**Figure 4E,F**). Iron loading decreases both *Tfrc* and *Slc11a2* expression in the liver during infection (**Figure 4C,E**). IAV infection reduces liver *Ftl1* expression in both the absence nd presence of iron loading, but significantly increases lung *Ftl1* expression in the presence of iron loading (**Figure 4G,H**). IAV infection increases liver and lung *Fth1* expression during iron loading (**Figure 4I,J**), agreeing with increased intensity of iron loading in cells observed with infection (**Figure S3A-D**). IAV infection does not impact *Slc40a1* expression in either the liver or the lung at 3dpi in the absence of iron loading, however, iron loading increases liver *Slc40a1* expression and IAV infection reduced this iron loading mediated increase (**Figure 4K,L**). IAV infection decreases liver *Aco1* expression in mice fed a CD, but has no effect on lung *Aco1* expression (**Figure 4M,N**). IAV infection reduces liver *Ireb2* expression and increases lung *Ireb2* expression, in mice fed a CD (**Figure 4O,P**). Iron loading reduces both *Aco1* and *Ireb2* expression in the liver in both the absence and presence of IAV infection. IAV infection has no effect on liver or lung *Hamp* expression in either the absence or presence of iron loading (**Figure 4Q,R**). Iron loading increases liver *Hamp* expression in both the absence and presence of IAV infection.

**Table S1: Human gene target primer oligonucleotide sequences used to measure mRNA expression *via* qPCR.**

| Gene (symbol) | Transcription direction | Nucleotide sequence (5'-3') |
|---|---|---|
| *Actin beta (ACTB)* | Forward | CTGGCACCACACCTTCTA |
| | Reverse | GGTGGTGAAGCTGTAGCC |
| *Transferrin receptor-1 (TFRC)* | Forward | AGGAACCGAGTCTCCAGTGA |
| | Reverse | ATCAACTATGATCACCGAGT |
| *Divalent metal transporter-1/solute carrier family 11 member 1 (SLC11A2)* | Forward | GGTGTTGTGCTGGGATGTTA |
| | Reverse | AGTACATATTGATGGAACAG |
| *Ferritin light chain (FTL)* | Forward | CCATGAGCTCCCAGATTCGT |
| | Reverse | TTCCAGAGCCACATCATCGC |
| *Ferritin heavy chain (FTH1)* | Forward | CCAGAACTACCACCAGGACT |
| | Reverse | CACATCATCGCGGTCAAAGT |
| *Ferroportin/solute carrier family 40 member 1 (SLC40A1)* | Forward | CTGTGCCCATAATCTCTGTC |
| | Reverse | CCATTTATAATGCCTCTTTCAG |
| *Iron regulatory protein-1/aconitase-1 (ACO1)* | Forward | GCGATGGACGCCCCAAAAGC |
| | Reverse | AGGCAGAACATCATTGGTGCC |
| *Iron regulatory protein-2/iron responsive element binding protein-2 (IREB2)* | Forward | CGTGCAGTCGGAGGAACAC |
| | Reverse | TCGAAAATGGTAAGCGCCCA |
| *Hepcidin (HAMP)* | Forward | CTGTTTTCCCACAACAGACG |
| | Reverse | CAGCACATCCCACACTTTGA |
| *Absent in melanoma-2 (AIM2)* | Forward | ATGCTGAATCTGACCAAAAG |
| | Reverse | GTTACCTTCTGGACTGGACTACAAAC |
| *NLR family CARD domain containing 4 (NLRC4)* | Forward | GAGGTCCCACAACTCGTCAA |
| | Reverse | GATTTCCCGCCAAATTCAAC |
| *NLR family pyrin domain containing 3 (NLRP3)* | Forward | GACTCTTGCACCCCGACTG |
| | Reverse | CTGGCTGGAGGTCAGAAGTGT |
| *Caspase-1 (CASP1)* | Forward | CTTCAGGCCACAGCAGAGCCCAAG |
| | Reverse | ACACACTTGAGGTCCCGGAGGA |
| *Caspase-4 (CASP4)* | Forward | GTGGTCCAGCCTCCATATTC |
| | Reverse | CGGGTCATGGCAGACTCTAT |
| *Interleukin-1 beta (IL1B)* | Forward | GAAGCTGATGGCCCTAAACA |
| | Reverse | AAGCCCTTGCTGTAGTGGTG |
| *Interleukin-18 (IL18)* | Forward | CAGTCTACACAGCTTCGGGA |
| | Reverse | TGCCACAAAGTTGATGCAAT |

**Table S2: Mouse gene target primer oligonucleotide sequences used to measure mRNA expression *via* qPCR.**

| **Gene (symbol)** | **Transcription direction** | **Nucleotide sequence (5'-3')** |
|---|---|---|
| Hypoxanthine phosphoribosyltransferase-1 *(Hprt1)* | Forward | AGGCCAGACTTTGTTGGATTTGAA |
| | Reverse | CAACTTGCGCTCATCTTAGGCTTT |
| *Transferrin receptor-1 (Tfrc)* | Forward | CCCATGACGTTGAATTGAACCT |
| | Reverse | GTAGTCTCCACGAGCGGAATA |
| *Divalent metal transporter-1/solute carrier family 11 member 2 (Slc11a2)* | Forward | AGCTAGGGCATGTGGCACTCT |
| | Reverse | ATGTTGCCACCGCTGGTATC |
| *Ferritin light chain (Ftl1)* | Forward | CGGAGGGTCAACATGCTATAA |
| | Reverse | AAGAGACGGTGCAGACTGGT |
| *Ferritin heavy chain (Fth1)* | Forward | GCTGAATGCAATGGAGTGTG |
| | Reverse | CAGGGTGTGCTTGTCAAAGA |
| *Ferroportin/solute carrier family 40 member 1 (Slc40a1)* | Forward | TGTCAGCCTGCTGTTTGCAGGA |
| | Reverse | TCTTGCAGCAACTGTGTCACCG |
| *Iron regulatory protein-1/aconitase-1 (Aco1)* | Forward | TGAAGGCCGAGTCCATCCTA |
| | Reverse | CGTTCACTCCCAAAGGCTCT |
| *Iron regulatory protein-2/iron responsive element binding protein-2 (Ireb2)* | Forward | ATGTCGAGGCCAGACTACCT |
| | Reverse | CTCAGGTTCAGGCACTGGTT |
| *Hepcidin (Hamp)* | Forward | CAACTTCCCCATCTGCATCTTC |
| | Reverse | GGGTGTAGAGAGGTCAGGATGTG |
| *Absent in melanoma-2 (Aim2)* | Forward | CTTCAGGCCACAGCAGAGCCCAAG |
| | Reverse | ACACACTTGAGGTCCCGGAGGA |
| *NLR family CARD domain containing 3 (Nlrc4)* | Forward | GAAAAGGATGGGAATGAAGCTC |
| | Reverse | TCCCCTCCAACTGCTTCAAC |
| *NLR family pyrin domain containing 3 (Nlrp3)* | Forward | GCTCCAACCATTCTCTGACC |
| | Reverse | AAGTAAGGCCGGAATTCACC |
| *Caspase-1 (Casp1)* | Forward | CTTCAGGCCACAGCAGAGCCCAAG |
| | Reverse | ACACACTTGAGGTCCCGGAGGA |
| *Caspase-4 (Casp4)* | Forward | TCATTTTACTCTGTCAAGCTGTCT |
| | Reverse | TGTCAGGGTGTTTGTTTTCAGC |
| *Interleukin-1 beta (Il1β)* | Forward | TGGGATCCTCTCCAGCCAAGC |
| | Reverse | AGCCCTTCATCTTTTGGGGTCCG |
| *Interleukin-18 (Il18)* | Forward | CGCCTCAAACCTTCCAAAT |
| | Reverse | GCCAAAGTTGTCTGATTCCA |
| *Interferon alpha-1 (Ifna1)* | Forward | ACCAACAGATCCAGAAGGCTCAAG |
| | Reverse | AGTCTTCCTGGGTCAGAGGAGGTT |

| | | |
|---|---|---|
| *Interferon alpha-4 (Ifna4)* | Forward | ACCAACAGATCCAGAAGGCTCAAG |
| | Reverse | AGTCTTCCTGGGTCAGAGGAGGTT |

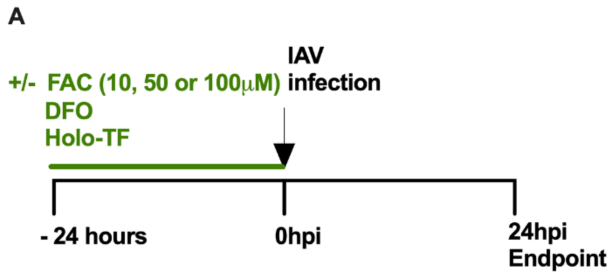


**Figure S1: Characterization and modulation of iron metabolism in human airway epithelial cell (AEC) models of influenza A virus (IAV) infection.** AECs (BCi-Ns1.1) were cultured with 10, 50 or 100µM of **A)** ferric ammonium citrate (FAC), deferoxamine (DFO) or holo-transferrin (Holo-TF) 24 hours prior to infection with A/H1N1/Auckland/1/2009 (H1N1) (MOI 0.5). Cells and supernatant were collected at **A)** 24 hours post-infection (hpi) or **B)** 6 and 12hpi.

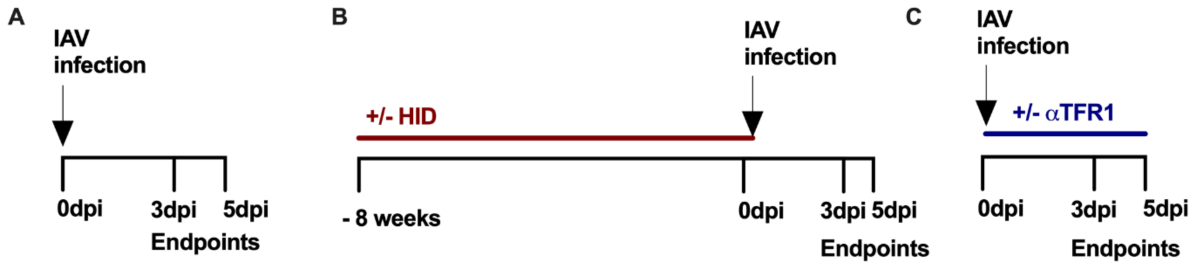


**Figure S2: Characterization and modulation of iron metabolism in murine models of influenza A virus (IAV) infection. A)** Six to eight week-old female BALB/c mice were intranasally inoculated with IAV (A/Puerto Rico/8/1934-H1N1 strain; 7.5PFU) on day 0 or sham-infected with media. Tissues were collected for assessing iron metabolism, infection and infection-induced disease at three or five days post-infection (dpi). **B)** Systemic and lung iron loading was induced in six to eight week-old female BALB/c mice by *ad libitum* consumption of high iron diet (HID; 2% carbonyl iron; 20g Fe/kg diet) for eight weeks. Control mice were fed a control diet (CD). Mice were intranasally inoculated with IAV (A/Puerto Rico/8/1934-H1N1 strain; 7.5PFU) on day 0 or sham-infected. Tissues were collected for assessing iron metabolism, infection and infection-induced disease at three or five dpi. **C)** Six to eight week-old female BALB/c mice were intranasally inoculated with IAV (A/Puerto Rico/8/1934-H1N1 strain; 7.5PFU) on day 0 or sham-infected. Mice were intranasally administered anti-transferrin receptor-1 monoclonal antibody (αTFR1; 1mg/kg) or isotype control (Iso; 1mg/kg) on day 0, 2 and 4. Tissues were collected for assessing iron metabolism, infection and infection-induced disease at three and five dpi.

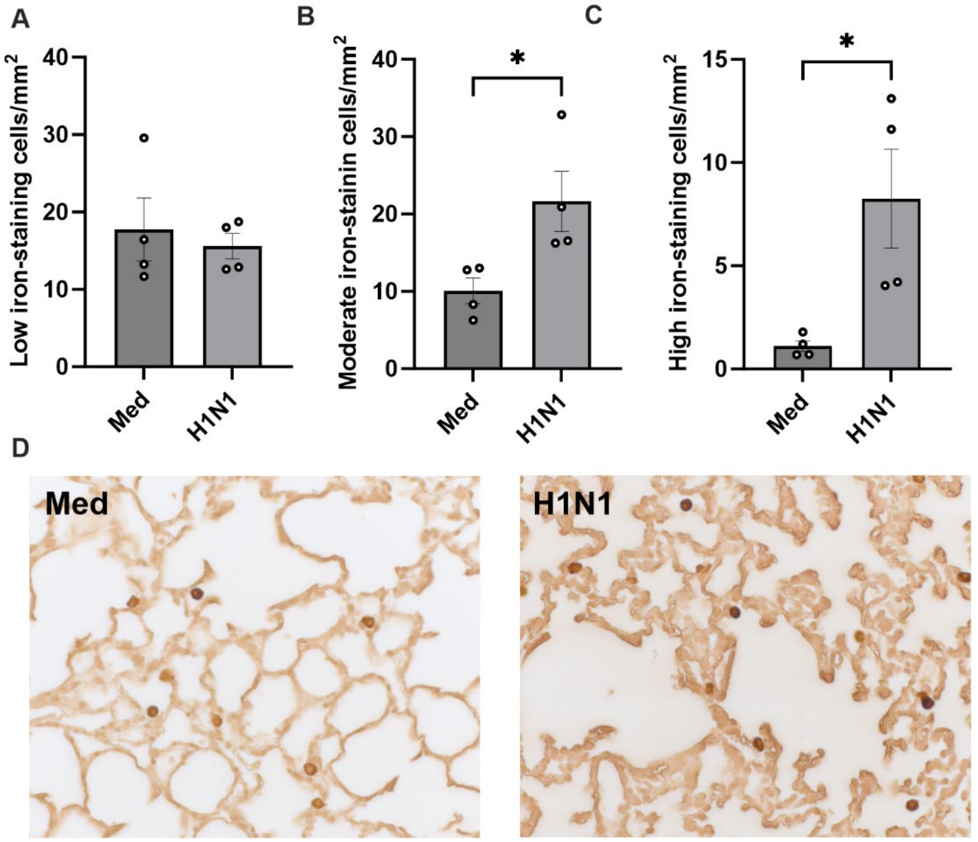


**Figure S3: Influenza A virus (IAV) infection increases cellular iron loading in lung tissues.** Mice were fed a control diet *ad libitum* for eight weeks prior to intranasal infection with IAV (H1N1) (A/PR8 strain; 7.5PFU) on day 0. At 5 days post-infection (dpi), lung histology sections were prepared and stained for iron using DAB-enhanced Perls staining. The numbers of cells that were positive for **A)** low, **B)** intermediate and **C)** high levels of iron staining per $mm^2$ of lung tissue were enumerated compared to cells in tissue sections from sham-infected controls. **D)** representative images of stained sections were also captured. Data (n=4) are presented as mean ± SEM *$p$<0.05; **$p$<0.01; ***$p$<0.001; ****$p$<0.0001.

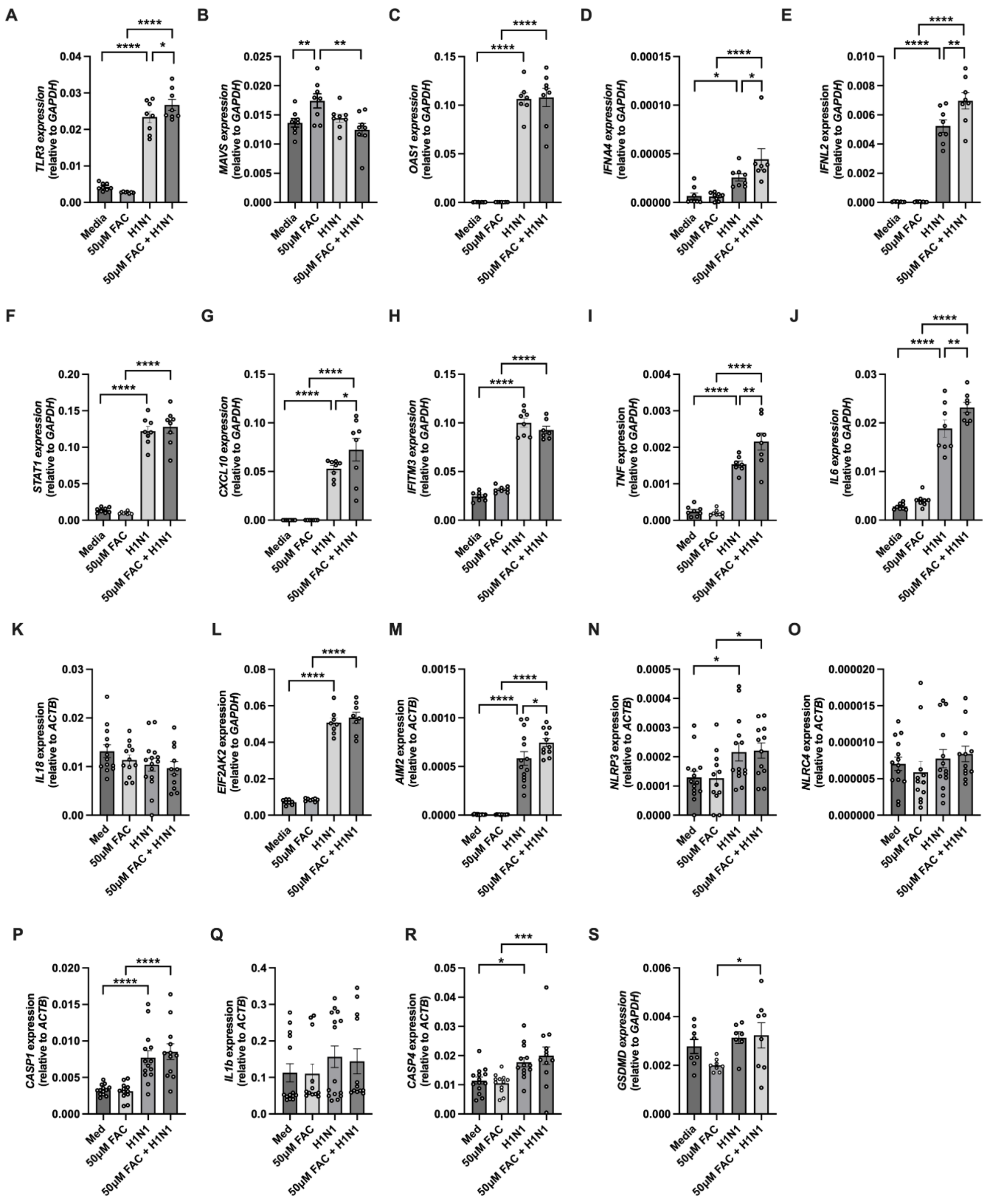


**Figure S4: Effect of increased iron availability on innate factor gene expression during influenza A virus (IAV) infection of human airway epithelial cells (AEC).** BCi-Ns1.1 cells were cultured with ferric ammonium citrate (FAC; 50µM) for 24 hours prior to infection with H1N1 (A/Auckland/1/2009) or sham-infected with media (Med). At 24 hours post infection, gene expression was assessed in AECs by qPCR for **A)** toll-like receptor 3 (*Tlr3*), **B)**

mitochondrial antiviral signalling protein (*Mavs*), **C)** 2'-5' oligoadenylate synthetase 1 (*Oas1*), **D)** interferon alpha 4 (*Ifna4*), **E)** interferon lamba 2 (*Ifnl2*), **F)** signal transducer and activator of transcription 1 (*Stat1*), **G)** CXC motif ligand chemokine 10 (*Cxcl10*), **H)** interferon induced transmembrane protein 3 (*Ifitm3*), **I)** tumour necrosis factor (*Tnf*), **J)** interleukin 6 (*Il6*), **K)** interleukin 18 (*Il18*), **L)** eukaryotic translation initiation factor 2 alpha kinase 2 (*Eif2ak2*), **M)** absent in melanoma 2 (*Aim2*), **N)** NLR family pyrin domain containing 3 (*Nlrp3*), **O)** NLR family CARD domain containing 4 (*Nlrc4*), **P)** caspase 1 (*Casp1*), **Q)** interleukin 1 beta (*Il1b*), **R)** caspase 4 (*Casp4*) and **S)** gasdermin D (*Gsdmd*)**.** Data (n=7-16) are presented as mean ± SEM. *$p<0.05$, **$p<0.01$, ***$p<0.001$, and ****$p<0.0001$.

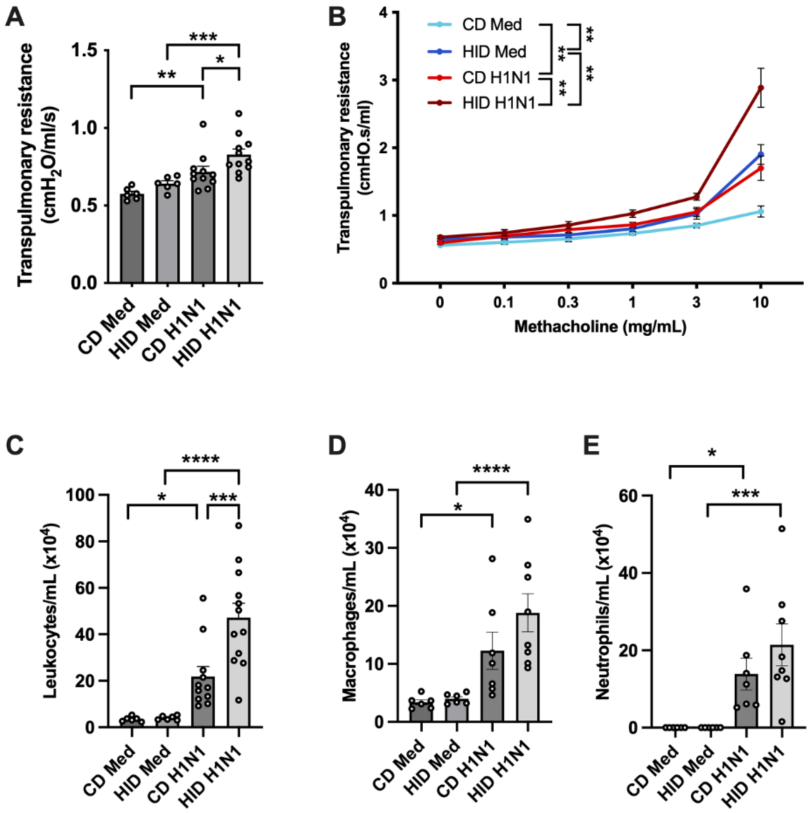


**Figure S5: Increased iron loading increases influenza A virus (IAV) infection-induced disease.** Mice were fed high iron diet (HID) or control diet (CD) *ad libitum* for eight weeks prior to intranasal inoculation with IAV (H1N1) (A/PR8 strain; 7.5PFU) on day 0. At 3 days post-infection (dpi), mice were assessed for lung function in terms of transpulmonary resistance at **A)** baseline and **B)** during challenge with increasing doses of nebulized methacholine and **C)** total leukocytes, **D)** macrophages and **E)** neutrophils in bronchoalveolar lavage fluid (BALF). Data (n=6-12) are presented as mean ± SEM. *$p$<0.05; **$p$<0.01; ***$p$<0.001; ****$p$<0.0001. Statistical significance is shown at 10mg/mL methacholine dose in **(B)**.

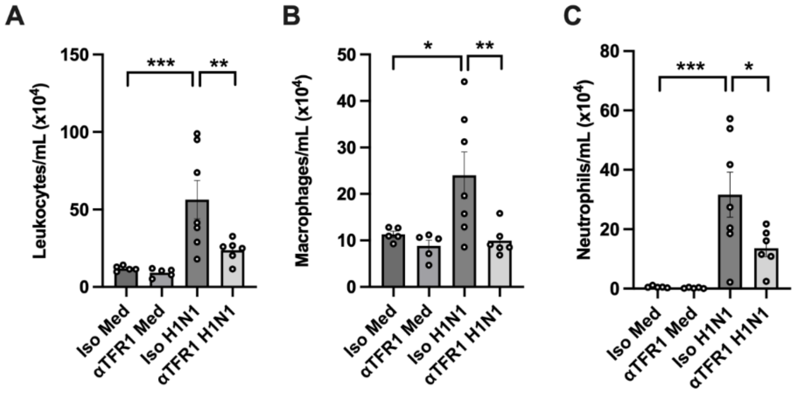


**Figure S6: Inhibition of TFR1 responses protects against influenza A virus (IAV) infection-induced inflammation**. Mice were intranasally infected with influenza A (H1N1) (A/PR8 strain; 7.5PFU) on day 0 and intranasally administered anti-transferrin receptor-1 monoclonal antibody (aTFR1; 1mg/kg) or isotype control (Iso; 1mg/kg) on day 0 and 2. At 3 days post-infection (dpi), mice were assessed for **A)** total leukocyte, **B)** macrophage and **C)** neutrophil numbers in bronchoalveolar lavage fluid (BALF). Data (n=5-7) are presented as mean ± SEM. **p*<0.05; ***p*<0.01; ****p*<0.001; *****p*<0.0001. Statistical significance is shown at 10mg/mL methacholine dose in **(B)**.

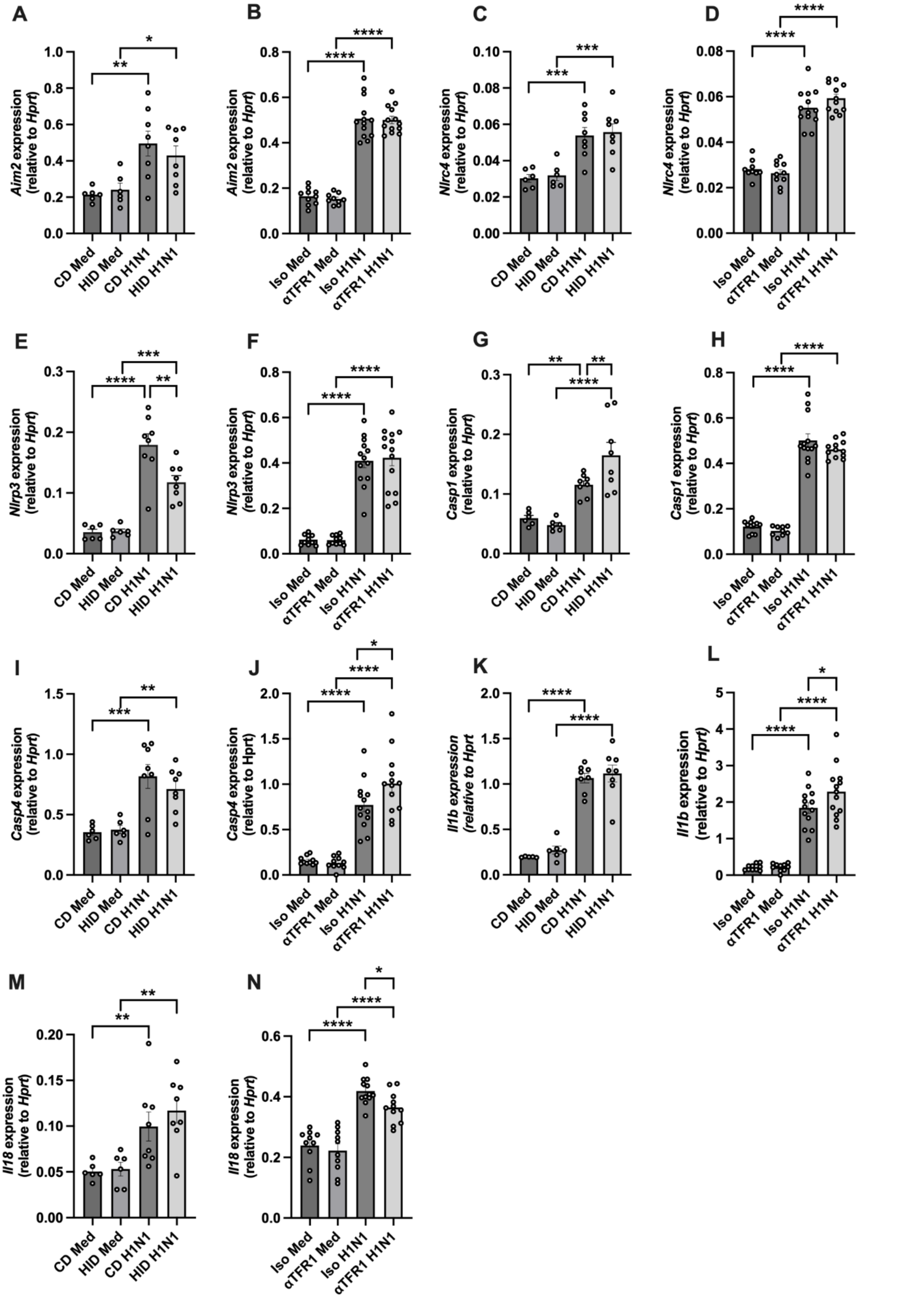


**Figure S7: Effects of increased iron loading, or inhibition of TFR1 responses, on specific inflammasome-related gene expression at five days post influenza A virus (IAV) infection.**

Mice were fed a high iron diet (HID) or control diet (CD) *ad libitum* for eight weeks prior to intranasal infection with H1N1 (A/Puerto Rico/8/1934) or sham-infected with media (Med). A separate group of mice were intranasally infected with H1N1 or sham-infected (d0) and intranasally administered anti-transferrin receptor-1 (αTFR1; 1mg/kg) or isotype control (Iso; 1mg/kg) monoclonal antibody on days 0, 2 and 4. At 5 days post-infection (dpi), gene expression was assessed in lung tissue by qPCR for **A, B)** absent in melanoma-2 (*Aim2*), **C, D)** NLR family CARD domain containing 4 (*Nlrc4*), **E, F)** NLR family pyrin domain containing 3 (*Nrlp3*), **G, H)** caspase-1 (*Casp1*), **I, J)** caspase-4 (*Casp4*), **K, L)** interleukin-1 beta (*Il1b)*, and **M, N)** interleukin-18 (*Il18*). Data (n=6-12) are presented as mean ± SEM. *$p<0.05$; **$p<0.01$; ***$p<0.001$; ****$p<0.0001$).

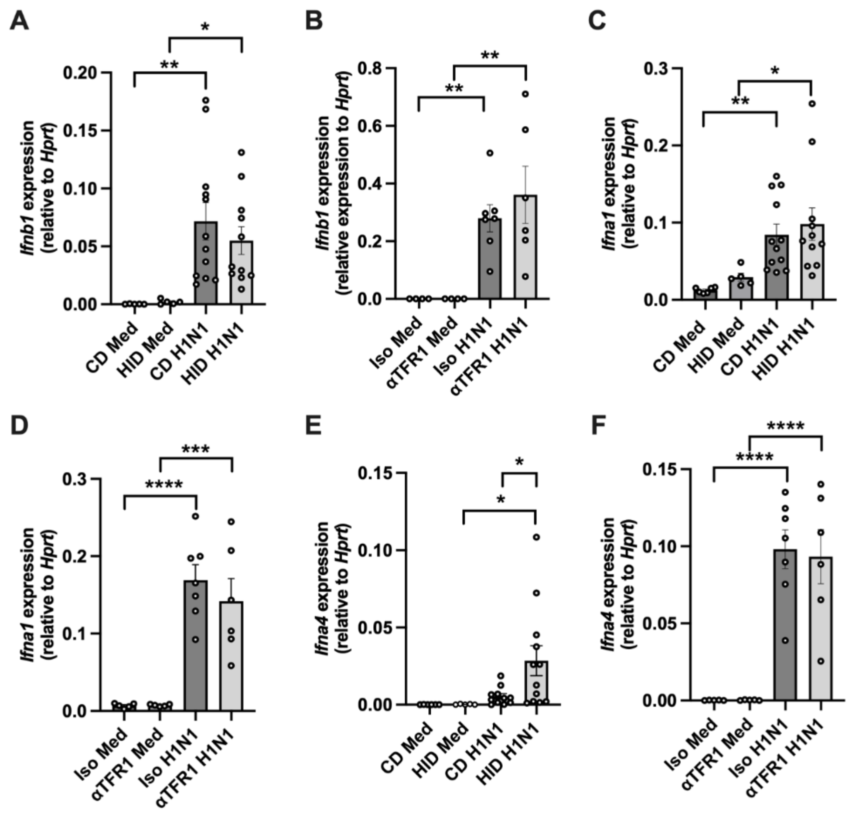


**Figure S8: Effect of increased iron loading, or inhibition of TFR1 responses, on type 1 interferon expression at three days post influenza A virus (IAV) infection.** Mice were fed a high iron diet (HID) or control diet (CD) *ad libitum* for eight weeks prior to intranasal infection with H1N1 (A/Puerto Rico/8/1934) or sham-infected with media (Med). A separate group of mice were intranasally infected with H1N1 or sham-infected (d0) and intranasally administered anti-transferrin receptor-1 (αTFR1; 1mg/kg) or isotype control (Iso; 1mg/kg) monoclonal antibody on days 0 and 2. At 3 days post-infection (dpi), gene expression was assessed in lung tissue by qPCR for **A, B)** interferon beta (*Ifnb*); **C, D)** Interferon alpha-1 (*Ifna1)* and **E, F)** Interferon alpha-4 (*Ifna4)***.** Data (n=5-12) are presented as mean ± SEM. *$p$<0.05; **$p$<0.01; ***$p$<0.001; ****$p$<0.0001.

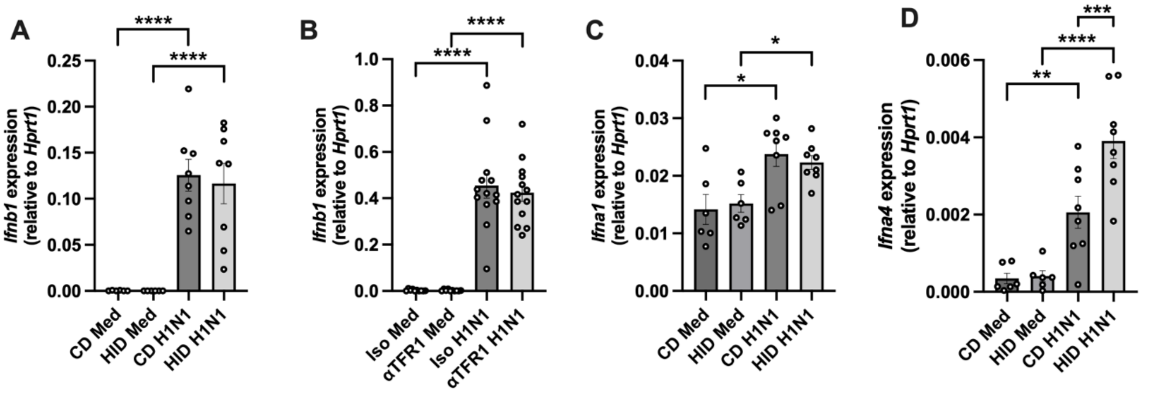


**Figure S9: Effect of increased iron loading, or inhibition of TFR1 responses, on type 1 interferon expression at five days post influenza A virus (IAV) infection.** Mice were fed a high iron diet (HID) or control diet (CD) *ad libitum* for eight weeks prior to intranasal infection with H1N1 (A/Puerto Rico/8/1934) or sham-infected with media (Med). A separate group of mice were intranasally infected with H1N1 or sham-infected (d0) and intranasally administered anti-transferrin receptor-1 (αTFR1; 1mg/kg) or isotype control (Iso; 1mg/kg) monoclonal antibody on days 0, 2 and 4. At 5 days post-infection (dpi), gene expression was assessed in lung tissue by qPCR for **A, B)** interferon beta (*Ifnb*); **C)** Interferon alpha-1 (*Ifna1)* and **D)** Interferon alpha-4 (*Ifna4)***.** Data (6-14) are presented as mean ± SEM. **p*<0.05, ***p*<0.01, ****p*<0.001, and *****p*<0.0001.
